# Lineage reversion drives WNT independence in intestinal cancer

**DOI:** 10.1101/2020.01.22.914689

**Authors:** Teng Han, Sukanya Goswami, Yang Hu, Fanying Tang, Maria Paz Zafra, Charles Murphy, Zhen Cao, John T Poirier, Ekta Khurana, Olivier Elemento, Jaclyn F. Hechtman, Rona Yaeger, Lukas E. Dow

## Abstract

The WNT pathway is a fundamental regulator of intestinal homeostasis and hyperactivation of WNT signaling is the major oncogenic driver in colorectal cancer (CRC). To date, there are no described mechanisms that bypass WNT dependence in intestinal tumors. Here, we show that while WNT suppression blocks tumor growth in most organoid and in vivo CRC models, the accumulation of CRC-associated genetic alterations enables drug resistance and WNT-independent growth. In intestinal epithelial cells harboring mutations in KRAS or BRAF, together with disruption of p53 and SMAD4, transient TGFβ exposure drives YAP/ TAZ-dependent transcriptional reprogramming and lineage reversion. Acquisition of embryonic intestinal identity is accompanied by a permanent loss of adult intestinal lineages, and long-term WNT-independent growth. This work delineates genetic and microenvironmental factors that drive WNT inhibitor resistance, identifies a new mechanism for WNT-independent CRC growth and reveals how integration of associated genetic alterations and extracellular signals can overcome lineage-dependent oncogenic programs.

---

The WNT signaling pathway is a key developmental regulator and essential for homeostasis of numerous adult cell types, including hematopoietic (1,2) and intestinal stem cells (3,4). Genetic alterations that hyperactivate the WNT pathway, including disruption of the APC tumor suppressor, mutational activation of CTNNB1 (β-catenin), or chromosomal translocations involving RSPO genes, occur in more than 90% of colorectal cancer (CRC) (5,6) and likely facilitate the initial growth of these tumors (7-9). We and others have shown that in many contexts, WNT hyperactivation is essential for tumor maintenance. For instance, restoration of endogenous APC expression in genetically engineered or tumor-engrafted mice leads to rapid and sustained tumor regression (10,11), while blocking WNT ligand secretion with porcupine (PORCN) inhibitors or blocking RSPO3 directly in human or murine tumors with RSPO3 fusions drives cell cycle arrest, differentiation, and tumor clearance (9,12,13). Thus, most normal and transformed cells of the intestine are thought to be WNT dependent.

Targeting the WNT pathway in cancer has clear therapeutic potential, and this has led to the recent initiation of multiple Phase I clinical trials (NCT02521844, NCT02649530, NCT03447470) for treatment of RSPO fusion cancers with PORCN inhibitors. However, due to the current lack of clinically approved WNT antagonists and scarcity of RSPO fusion model systems, it has not been possible to explore potential mechanism of therapy failure and resistance to WNT-targeted therapy. To further understand WNT dependence and the therapeutic potential of WNT inhibition, we engineered an array of organoid-based models capturing the most frequently observed genetic alterations in this molecular subtype. We found that while no individual oncogenic change influences sensitivity to WNT suppression, the combined alteration of multiple CRC-associated lesions enables rapid acquired resistance to WNT blockade. This occurs via a TGFβ-induced lineage conversion that reverts adult intestinal epithelium to a fetal intestinal state. The lineage switch is driven and maintained by YAP/TAZ signaling, and is not reversible. Consequently, WNT-independent cells become exquisitely and selectively sensitive to genetic or pharmacologic suppression of YAP/TAZ.

## RESULTS

### Accumulation of CRC-associated oncogenic mutations leads to WNT independence

PTPRK-RSPO3 fusions drive the development of WNT-dependent murine intestinal adenomas that are extremely sensitive to treatment with the PORCN inhibitor WNT974 (9). Human RSPO3 fusion tumors carry an array of cooperating oncogenic lesions, including frequent mutational activation of KRAS or BRAF, as well as disruption of TP53 and/or SMAD4 (Figure 1a). How these co-occurring alterations contribute to WNT-dependence is not known. Unlike APC mutant CRC, there are very few human RSPO3 cell lines or organoid models with which to interrogate RSPO disease biology. To recapitulate some of the genetic complexity of human RSPO3 fusion tumors, we used CRISPR to develop a series of murine organoid models harboring common oncogenic genotypes (Figure 1b). We first generated multiple independent organoid lines, each carrying an endogenous Ptprk-Rspo3 (R) fusion (9), and an endogenous *Kras*^*G12D*^ (K) mutation (from the *LSL-Kras*^*G12D*^ allele) (14). We then used sgRNAs targeting *Trp53* (P) and *Smad4* (S) to create triple (KRP and KRS) and quadruple (KRPS) mutants (Figure 1b-c; see methods for details). As previously described (15,16), organoids containing loss of function alterations in p53 and SMAD4 were selected by treating for seven days with Nutlin-3 (10μM) or TGFβ (5ng/ ml), respectively. We confirmed the presence of each mutation by sequencing or assessed protein disruption by western blot (Figure 1d-f).

**Figure 1.**
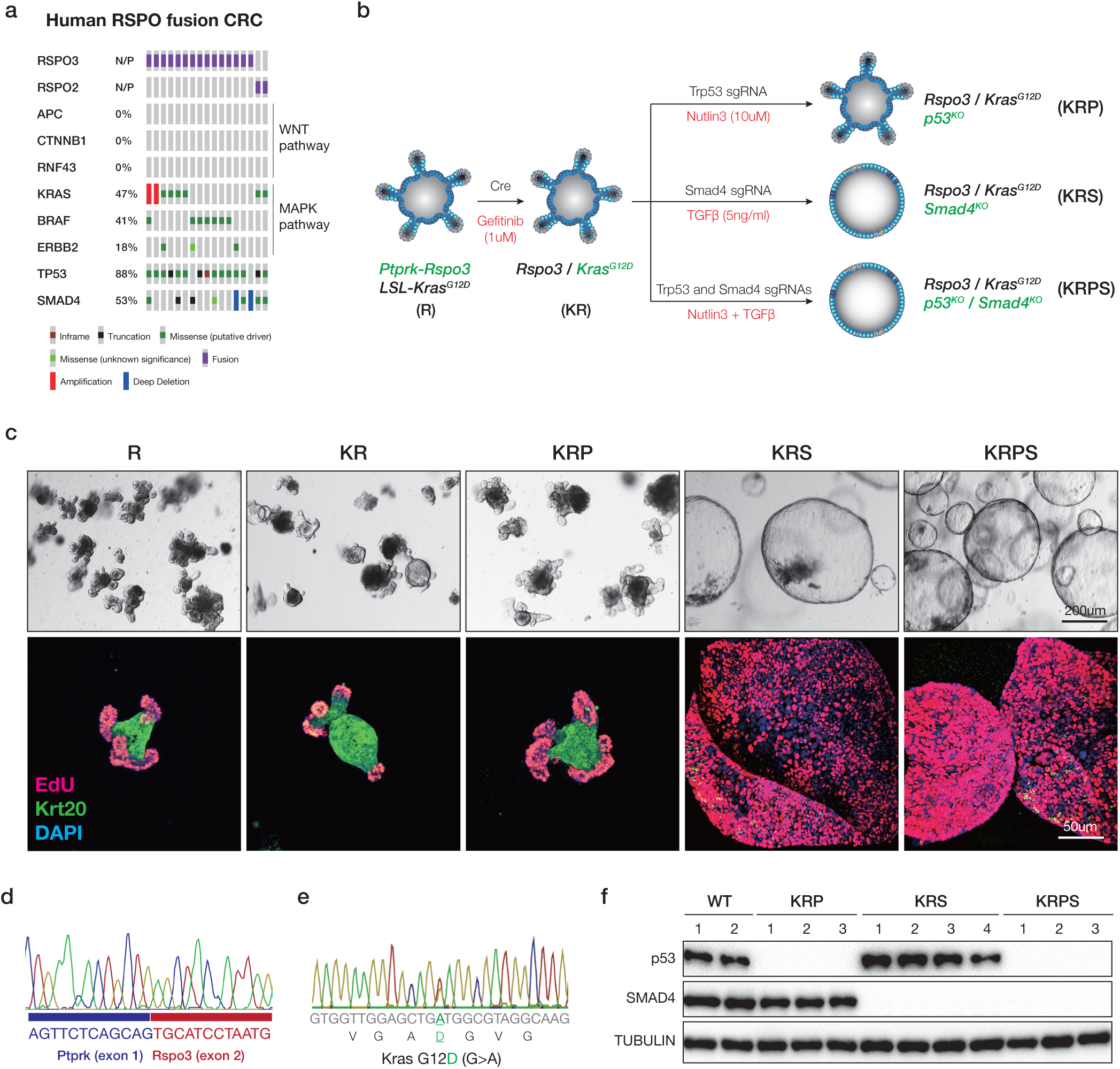
Engineering of mouse-derived intestinal tumour organoids. **a.** Oncoprint showing co-ocurring mutations in WNT, MAPK, TP53 and SMAD4 genes in R-Spondin fusion-positive human CRCs. b. Schematic representation of the generation of sequentially mutated murine intestinal organoids. c. Bright field (upper) and immunofluorescent (lower) images of sequential mutants. Proliferating cells are marked by EdU (Red), and differentiated cells by KRT20 (green). d. Representative sanger sequencing chromatogram confirming expected Ptprk-Rspo3 mRNA fusion junction following CRISPR-mediated intrachromosomal inversion. e. Representative Sanger sequencing chromatogram from cDNA, confirming expression of the KrasG12D (G>A) mutant transcript. f. Western blots showing loss of p53 and SMAD4 in independent biological replicates of edited and selected organoids.

Similar to organoids carrying only Rspo3 fusions (9), KRP and KRS organoids showed rapid cell cycle arrest (loss of EdU incorporation) and differentiation (induction of KRT20) 4 days following PORCN inhibition (Figure 2a-b). Quadruple KRPS mutants showed a marked decrease in EdU incorporation and increased KRT20 expression, however, unlike KRP and KRS, a subpopulation of KRPS cells (4-5%) continued to proliferate and expand in the presence of drug (Figure 2a-b). Resistant cells maintained very low levels of WNT target genes, suggesting they did not escape WNT974 treatment by downstream activation of WNT signaling (Figure 2c). This was not specific to WNT974 as we observed an identical response with an independent PORCN inhibitor (ETC-159) (Figure 2b). Resistance to both drugs was long-lived as re-challenge after 2-4 weeks in culture caused only a minor reduction in proliferation (Figure 2c) and no change in cell morphology (Supplementary Figure 1a). Further, quadruple mutant organoids expressing an endogenous *Braf*^*V619E*^ allele (“BRPS”) also showed rapid resistant outgrowth in the presence of WNT974 (Supplementary Figure 1b).

**Figure 2.**
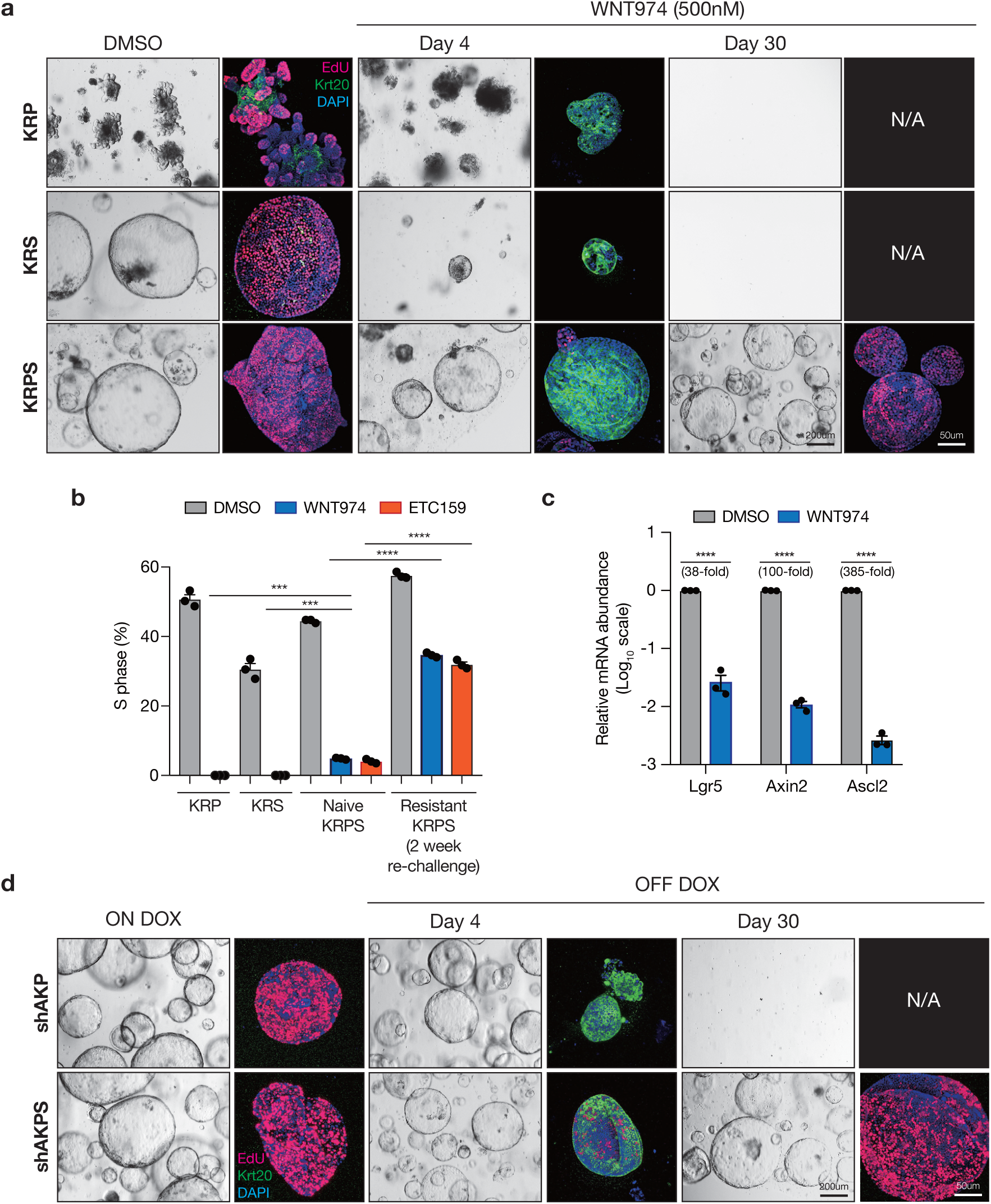
Kras, p53 and Smad4 mutations enable Wnt independence. **a.** Bright field and immunofluorescent images of KRP, KRS and KRPS organoids treated with DMSO or WNT974. Proliferating cells are marked by EdU (Red), and differentiated cells by KRT20 (green). **b**. EdU flow cytometry of KRP, KRS, naive KRPS and resistant KRPS organoids treated with DMSO or WNT974 (n=3 independent biological replicates per condition). **c.** qPCR of canonical WNT targets showing WNT signaling is not reactivated in WNT974-resistant KRPS organoids. **d**. Bright field and immunofluorescent images of shAKP and shAKPS organoids before and following APC restoration (dox withdrawal). EdU (red) and KRT20 (green) mark proliferating and differetiated cells, as above. For all graphs, error bars represent the +/- standard error of the mean (SEM).

To determine whether the (KRPS/BRPS) genotype-dependent response was specific to PORCN inhibition, or reflected a more general bypass of WNT-dependence, we generated organoids carrying KRAS^G12D^ and p53 mutations and an inducible Apc shRNA that enables doxycycline (dox)-regulated control of APC expression. In this context, withdrawal of dox drives APC-mediated suppression of hyperactive WNT signaling and in the absence of exogenous RSPO1/WNT ligand, promotes cell cycle arrest and differentiation (10). Consistent with our previously published work, APC restoration in shAKP cells induced cell cycle arrest and differentiation, preventing RSPO-independent growth (Figure 2d). However, similar to KRPS cells, a subpopulation of SMAD4 mutant ‘shAKPS’ cells were able to expand in RSPO-free media following APC re-expression (Figure 2d). Further, these RSPO/WNT-independent shAKPS cells showed no acute response to treatment with WNT974, confirming they did not escape APC restoration by upregulating WNT ligand expression (Supplementary Figure 1c). Together, these data suggest that the combination of *Kras/Braf, p53* and *Smad4* mutations can enable the generation of WNT-independent intestinal organoids, following pharmacologic or genetic suppression of WNT signaling.

In total, we generated 21 WNT-independent organoid lines, which showed two distinct mechanisms of resistance. Approximately one quarter (5/21) showed robust reactivation of the WNT pathway (Supplementary Table 1), in some instances via genomic amplification or hotspot mutation of β-catenin (Supplementary Figure 2a-c). Using base editing, we confirmed that mutational activation of *Ctnnb1* (S33F) was sufficient to enable WNT974 resistant outgrowth (Supplementary Figure 2d). The majority of WNT-independent organoids (16/21) showed no evidence of WNT activating mutations (of those examined by WES) and remained responsive to WNT-suppressive stimuli (Supplementary Figure 2e). This work reveals the unexpected appearance of a population of intestinal cells that remain responsive to WNT-suppressive stimuli, yet do not require WNT pathway activation for growth and proliferation. We defined organoids with this “resistant” phenotype as KRPS^R^, BRPS^R^ and shAKPS^R^.

### Transient TGFβ stimulation is required for resistance to WNT inhibition

Transcriptome analysis of WNT974-naïve and resistant isogenic organoid pairs by RNAseq revealed a number of differentially regulated genes and pathways in both KRPS^R^ and shAKPS^R^ genotypes, including a series of inflammation-associated gene signatures (Figure 3a). We noted that during the generation of KRPS, BRPS, and shAKPS cultures, each sample was treated transiently with TGFβ to select for SMAD4 mutant populations (Figure 1b) and reasoned that TGFβ-treatment might be altering transcriptional networks and conditioning or ‘priming’ cells to become WNT-independent. To determine whether TGFβ-priming was important for WNT independence, we generated KRPshS organoids in which SMAD4 expression was silenced by an shRNA, enabling the enrichment of cells with SMAD4 depletion, without functional (TGFβ) selection (Figure 3b). In contrast to spheroid KRPS cells, KRPshS cultures retained a budding organoid morphology similar to R, KR, or KRP cells (Figure 1c, Figure 3c). However, short-term treatment (3 days) with TGFβ (5ng/ml) induced a rapid morphological shift toward spheroid structures (Figure 3c-d) that was maintained indefinitely following withdrawal of TGFβ (Figure 3d). The spheroid change was due to signaling via the canonical TGFβ receptor complex as co-treatment with the TGFβR1 inhibitor (LY2157299) or CRISPR-mediated genetic disruption of *Tgfbr2* completely abolished the morphological response (Figure 3c-d).

**Figure 3.**
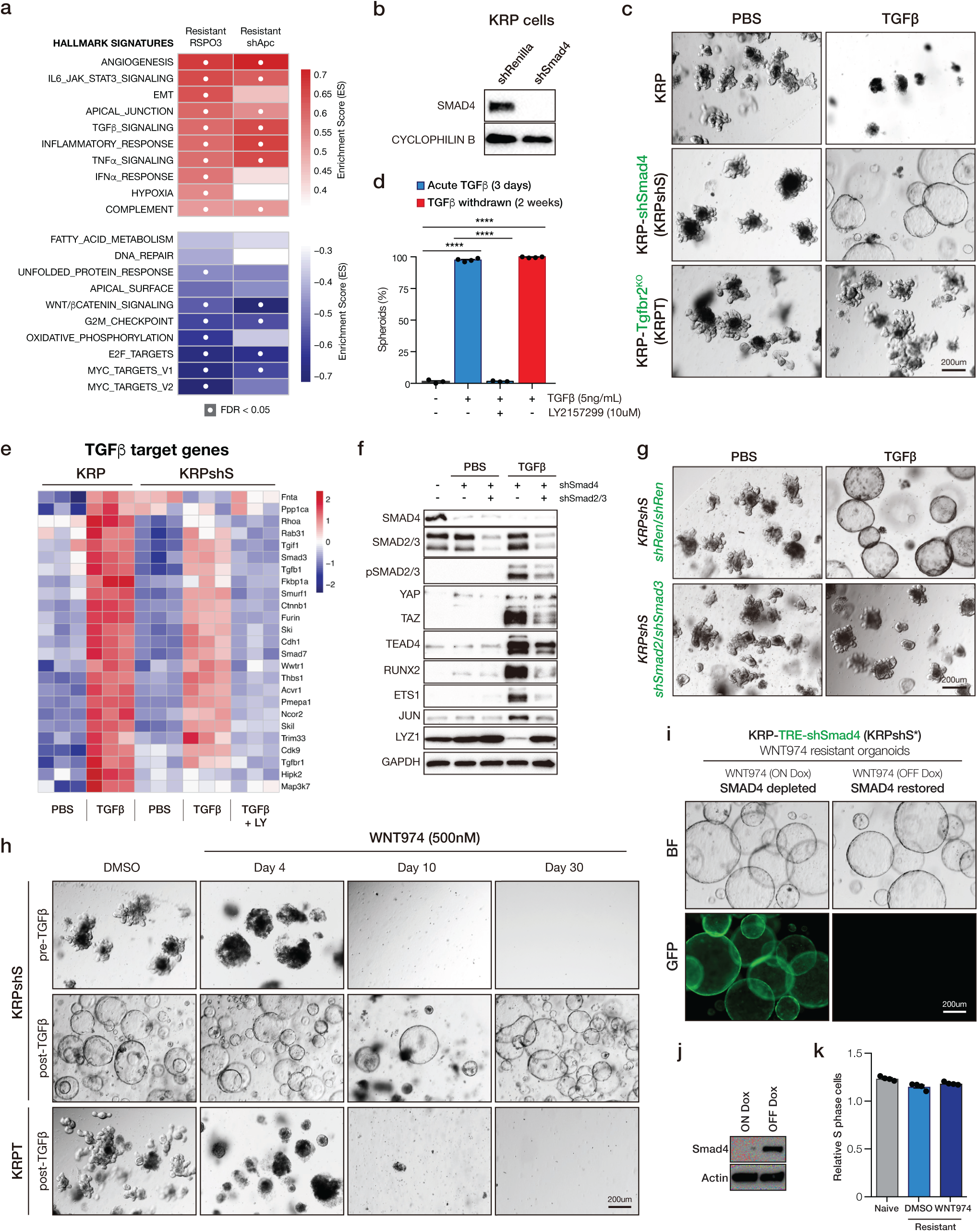
TGFβ is required for WNT independence. **a.** GSEAs show inflammation-related pathways are enriched in WNT independent lines. **b**. Western blot analysis showing Smad4 knockdown. **c**. Bright field images of KRP, KRPshS and KRPT organoids treated with PBS or 5ng/mL TGFβ as indicated. **d.** Quantification of TGFβ-induced organoid morphology change. **e.** Heatmap of TGFβ target genes in KRP and KRPshS acutely treated with TGFβ. **f.** Western blots showing Smad2/3 depletion inhibits the induction of TGFβ downstream targets. **g.** Bright field images showing that Smad2/3 depletion blocks TGFβ-induced spheroid formation. **h.** Bright field images of KRPshS (pre- and post-TGFβ) and KRPT (post-TGFβ) showing TGFβ priming is required for WNT974 resistance. **i.** Bright field images of Smad4 restoration in KRPS WNT974-resistant line showing Smad4 is not required for WNT independence. **j**. EdU flow cytometry on Smad4 restored WNT974-naive KRPS and WNT974-resistant KRPS lines. **k**. Western blot showing Smad4 is restored in WNT974-resistant KRPS. Error bars represent +/- standard error of the mean (SEM).

In many contexts, SMAD4 association with SMAD2/3 is required for canonical TGFβ signaling (17). Indeed, the magnitude of TGFβ target gene expression following acute TGFβ stimulation was reduced in SMAD4-depleted cells (KRPshS) compared to those with wildtype SMAD4 (KRP). However, most downstream targets were still strongly induced compared to untreated cells, highlighting a widespread SMAD4-independent transcriptional response to TGFβ in these cells (Figure 3e, Supplementary Figure 3a), as has been reported in other settings (18,19). In support of the notion that TGFβ drives SMAD4-independent canonical signaling, we saw a marked increase in SMAD2/3 phosphorylation, but no change in the activation of non-canonical TGFβ pathways (MAPK, Akt, p38, JNK) (Supplementary Figure 3b). Further, TGFβ-mediated transcriptional changes in SMAD4-depleted cells were SMAD2/3-dependent, as simultaneous silencing of these SMADs suppressed target gene induction and spheroid transformation (Figure 3f-g). Most importantly, and consistent with results from CRISPR-derived KRPS cultures, TGFβ-treated KRPshS organoids developed resistance to WNT974, while TGFβ-naïve KRPshS organoids remained WNT-dependent, and could not survive continuous treatment with the drug (Figure 3h). Thus, TGFβ priming initiates the induction of WNT-independence.

As noted above, withdrawal of TGFβ from the culture media had no impact on TGFβ-induced spheroid morphology (Figure 3d). Likewise, treatment of WNT-independent organoids with a TGFβR1 inhibitor (LY2157299) had no effect on morphology or growth of organoids (Supplementary Figure 3c-d). This observation raised the possibility that SMAD4 loss was important to enable TGFβ priming, but was dispensable beyond this event. To directly test the requirement for SMAD4 loss in regulating WNT-independence we generated KRPshS organoids in which the expression of the SMAD4 shRNA was controlled by a dox-regulated element (TRE) and a reverse tet-transactivator (Supplementary Figure 3e). We generated WNT974-resistant organoids in the presence of dox, exactly as described above, and restored SMAD4 expression by withdrawal of dox from the media (Figure 3i-j). As expected, re-expression of SMAD4 sensitized organoids to acute exposure to TGFβ (Supplementary Figure 3f-h), however it had no impact on organoid morphology, proliferation, or WNT independence in the absence of exogenous TGFβ ligand (Figure 3i,k). Together, these data support the idea that transient TGFβ-priming in KRPS/shAKPS intestinal cells drives SMAD2/3-dependent TGFβ pathway activation that avoids SMAD4-dependent cell death (19), but that continual TGFβ signaling is not essential to maintain the WNT independent state.

### The in vivo tumor microenvironment is sufficient to prime cells for WNT independence

TGFβ is an abundant cytokine in the tumor microenvironment and modulation of TGFβ receptor has been shown to impact colorectal cancer progression in animal models by altering tumor-stroma signaling (20,21). We sought to determine whether exposure to tumor microenvironmental signals were sufficient to prime TGFβ-naïve organoids and induce WNT independence in vivo.

Long-term treatment with WNT974 can cause intestinal and bone damage in mice (22,23). To enable potent and sustained WNT suppression in tumor cells without effects on surrounding tissues, we used the regulatable shApc model, whereby withdrawal of dox from chow drives rapid APC restoration and potent WNT pathway suppression (10,11). We generated TGFβ-naïve shAKPS cells by simultaneous delivery of sgRNAs targeting both Trp53 and Smad4, and selected for p53 disruption by Nutlin-3 treatment. Surrogate selection for p53 targeting produced a population of cells that were TGFβ-naïve, but contained greater than 95% Smad4 disruption (Supplementary Figure 4a-b). As expected, TGFβ-naïve shAKP and shAKPS organoids, and TGFβ-treated shAKPT organoids were sensitive to WNT suppression by APC restoration, and never became WNT-independent in vitro (Supplementary Figure 4c).

Each of the organoid lines were injected in mice pre-treated with dox (200mg/kg in chow). When tumors reached ∼100mm^3^, animals were either maintained on dox or returned to normal chow to restore APC. As previously described, shAKP organoids showed cell cycle arrest (loss of Ki67) and differentiation (increased KRT20) in response to APC restoration (Figure 4a), and showed little to no tumor growth following the dox switch (Figure 4b). As expected, tumors derived from TGFβ-treated shAKPS organoids continued to grow following APC restoration (Figure 4a). At harvest, these tumors were highly proliferative and showed little evidence of differentiation, similar to dox-treated tumors. Tumors derived from TGFβ-naïve shAKPS organoids, that were entirely WNT-dependent in vitro (Supplementary Figure 4c), showed a mixed response in vivo, with clusters of arrested differentiated cells interspersed with poorly differentiated and proliferative tumor cells (Figure 4a). This is consistent with the outgrowth of a subset of WNT independent tumor cells, as observed in TGFβ-primed ex vivo cultures (Figure 2b). Continued growth of shAKPS cells was most likely due to canonical TGFβ-mediated signaling, as genetic disruption of *Tgfbr2* did not allow WNT-independent tumor growth (Figure 4c-d). Similar effects were seen with KRPS-derived tumors, following 2 weeks of WNT974 treatment (Supplementary Figure 5). In all, these data show that the tumor microenvironment is sufficient to prime WNT-dependent cells to become WNT independent. Lineage reversion underlies WNT independence

**Figure 4.**
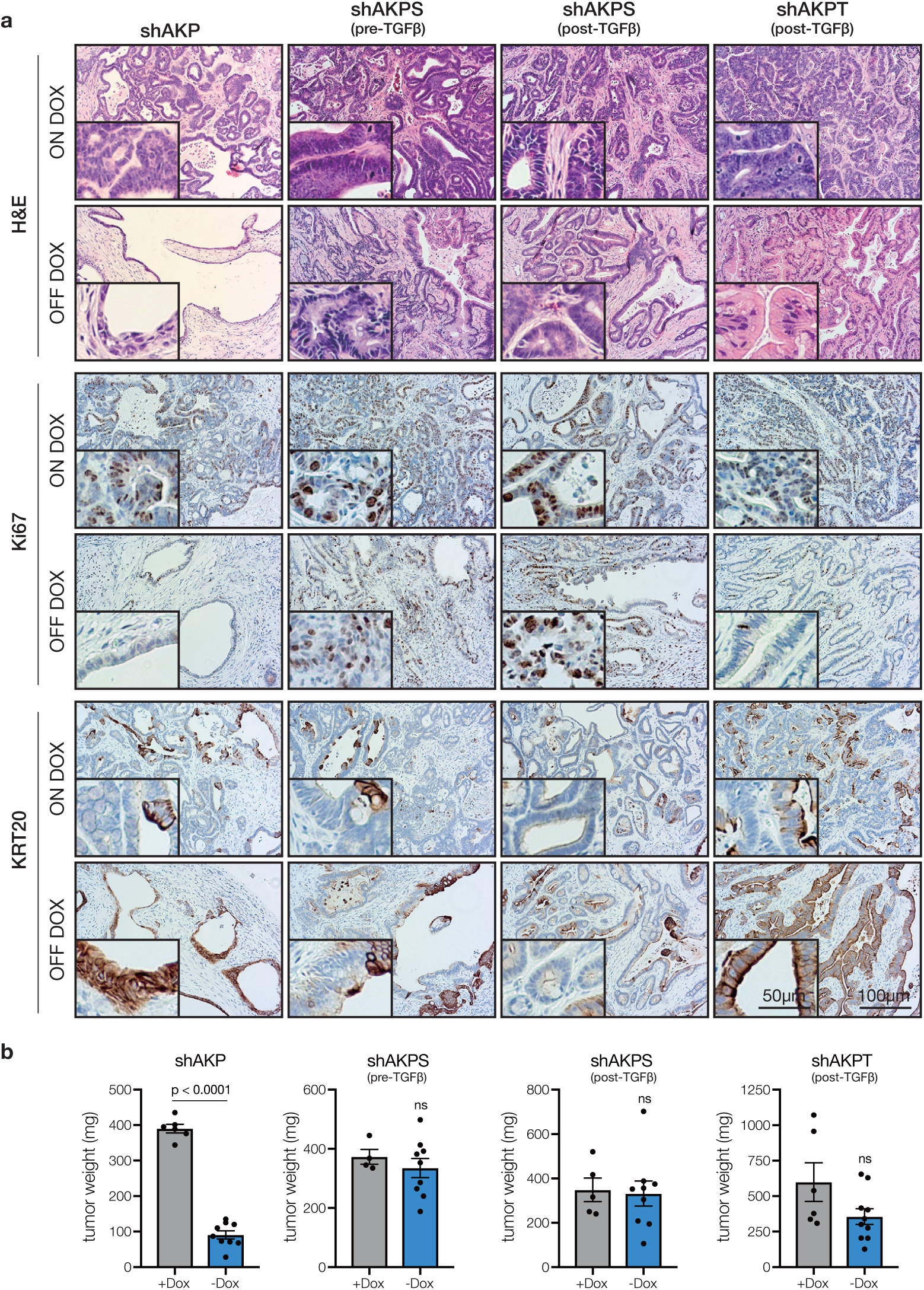
The in vivo tumor microenvironment is sufficient to prime cells for WNT independence. **a.** Immunohistochemical stains of organoid transplants on dox for 6 weeks (ON DOX), or on dox for 1 week then off dox for 5 weeks. **b.** Tumor weight quantification following transplant of different organoid genotypes. Error bars represent +/- standard error of the mean (SEM), n≥4 individual tumors, p-values calculated using a two-sided t-test, with Welsh’s correction. While shAKPT tumors showed dramatic molecular and histopathogical response, tumor volume was not statistically significantly different (p=0.14), likely owing to increased growth rate of this organoid genotype.

Given that WNT independence was initiated, but not maintained by TGFβ, we sought to identify downstream signaling or pathway alterations that were altered following TGFβ exposure and maintained following TGFβ withdrawal. To do this, we again compared pathway enrichment using the hallmark GSEA dataset (24), as well as a range of unique gene signatures that have recently been identified from single-cell and bulk transcriptome analyses of the mouse intestine (25-28). As expected, TGFβ signaling was the most enriched pathway following acute (3d) TGFβ treatment, but was lost after long-term (>30d) TGFβ-free culture (Figure 5a). Strikingly, the only molecular signature that was increased and robustly maintained in post-TGFβ organoids was a collection of genes upregulated in embryonic or fetal intestinal organoids, compared to adult-derived cultures (Figure 5a-b) (28). Accordingly, genes downregulated in fetal intestinal cultures were also strongly suppressed following TGFβ-priming (Figure 5a). Many of the genes downregulated in TGFβ-primed organoids specified differentiated intestinal lineages, including enterocytes, enteroendocrine, and Paneth cells (Figure 5a-b). These changes identified by transcriptome analysis were readily apparent in tumors in vivo, with KRPS and shAKPS tumors, but not KRP, shAKP, KRPT, or shAKPT tumors, showing loss of adult lineages (Paneth cells) and induction of fetal markers (SPP1 and ANXA1) (Figure 5c, Supplementary Figure 6)

**Figure 5.**
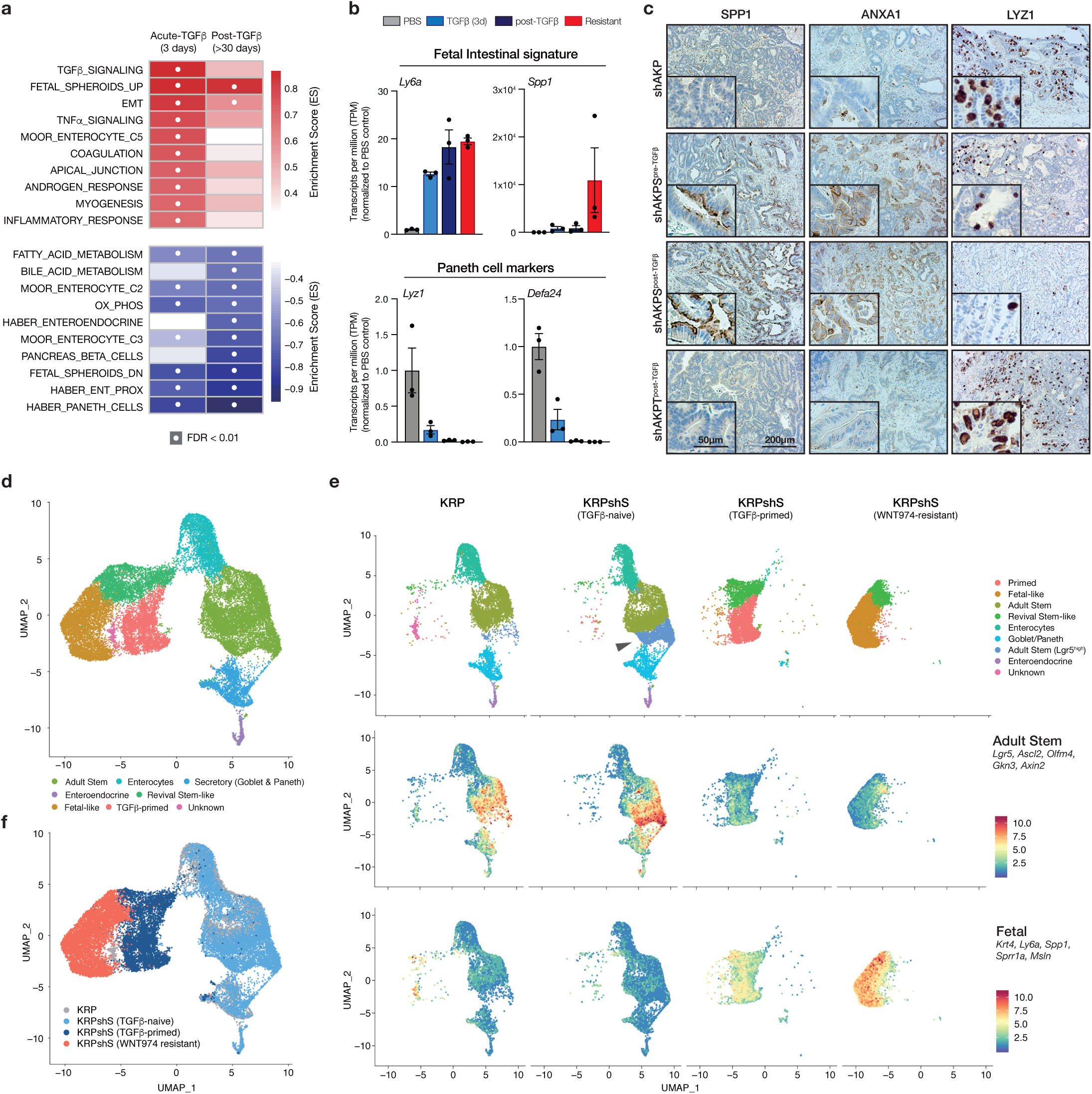
TGFβ drives fetal-like lineage reversion. **a.** GSEA summary showing pathway enrichment scores for KRPshSmad4 organoids treated for 3 days with TGFβ (acute) and the same organoids one month after TGFβ withdrawal. White dots represent significantly enriched gene sets (FDR<0.01). **b**. mRNA expression (transcript per million estimates) of fetal intestinal markers (upper) and Paneth cell markers (lower) in KRPshS organoids. Error bars are +/- SEM. **c.** Immunohistochemical staining of SPP1, ANXA1, and LYZ1 on organoid-derived tumors, as labelled. **d.** UMAP plot of merged scRNAseq from KRP, KRPshS (pre-TGFβ), KRPshS (post-TGFβ), and KRPshS (WNT974-resistant) organoids, showing identified and putatively identified cell populations. **e**. UMAP plot (upper) of individual scRNAseq samples shows expansion of Lgr5^high^ adult stem cell population following Smad4 silencing (arrow), and transition of cell lineage following TGFβ-priming and development of WNT-independence. Feature plots (middle and lower) highlight the mean expression of adult stem cell and fetal intestinal markers, as indicated. **f**. UMAP plot of merged scRNAseq data highlighting specific samples.

To more closely interrogate the dynamics of cell fate change following TGFβ priming and the development of WNT independence, we performed single cell RNA sequencing (scRNAseq) in KRP, and KRPshS TGFβ-naïve, TGFβ-primed, and WNT974-resistant organoids. Mapping of marker gene expression across 12 identified cell clusters revealed multiple distinct populations, including Lgr5-positive adult stem cells, enterocytes, goblet, Paneth, and enteroendocrine cells (Figure 5d; Supplementary Figure 7). Depletion of SMAD4 (in KRPshS organoids) induced expansion of a Wnt-high and Lgr5-high stem cell compartment compared to KRP (Figure 5e, Supplementary Figure 7, Supplementary Figure 8), as reported in other intestinal organoid and in vivo models (29,30). Despite the expansion of the stem cell pool, SMAD4 loss alone did not alter lineage differentiation (Figure 5e, upper; Supplementary Figure 7). In contrast, TGFβ priming of KRPshS cells caused a dramatic shift in transcriptional identity, depleting multiple differentiated intestinal lineages (Figure 5e-f). In addition, we noted a minor expansion of recently identified ‘revival stem cells’ (expressing *Clu, Anxa1, Cxadr*, and *Basp1*; hereafter “RevSCs”) that are activated in vivo following intestinal injury (31) (Supplementary Figure 8a-b). WNT974-resistant KRPshS cells were transcriptionally distinct from TGFβ-primed cultures, with some overlap in the putative RevSC cluster. They showed further depletion of differentiated intestinal lineages (Supplementary Figure 8c) as well as low Wnt and adult stem cell signatures (Figure 5f, middle; Supplementary Figure 7; Supplementary Figure 8b). Most notably, WNT974-resistant organoids showed marked elevation of the fetal intestinal signature (Figure 5e, lower), consistent with a role for this transition in the development of WNT-independence.

### YAP/TAZ signaling is necessary and sufficient to drive lineage reversion and WNT independence

YAP1 and TAZ (WWTR1) are closely related transcriptional co-activators in the Hippo signaling pathway that have been linked to the acquisition of stem and progenitor cell properties in breast, pancreas, and neuronal tissue (32), and for tissue regeneration in the colonic epithelium (33). YAP1 and TAZ are direct transcriptional targets of SMAD2/3 (34,35) (Supplementary Figure 9a), and were induced following TGFβ treatment (Figure 6a). YAP transcriptional signatures were strongly induced by TGFβ treatment and remained high following TGFβ-withdrawal (Figure 6a-b). Remarkably, the expression of a small set of recently reported canonical YAP target genes (36) was sufficient to accurately cluster the organoid genotypes into pre-TGFβ, post-TGFβ, and WNT-independent subtypes (Supplementary Figure 9b). Moreover, this comparison, along with single-cell analysis highlighted that the relative amplitude of the YAP signature was markedly increased in WNT-independent organoids relative to TGFβ-primed cells (Figure 6c, Supplementary Figure 9b). Finally, analysis of chromatin accessibility TGFβ-primed and WNT974 resistant organoids by ATACseq revealed the consensus TEAD binding motif as the most significantly enriched of all known transcription factor binding motifs in WNT-independent cells (Figure 6d).

**Figure 6.**
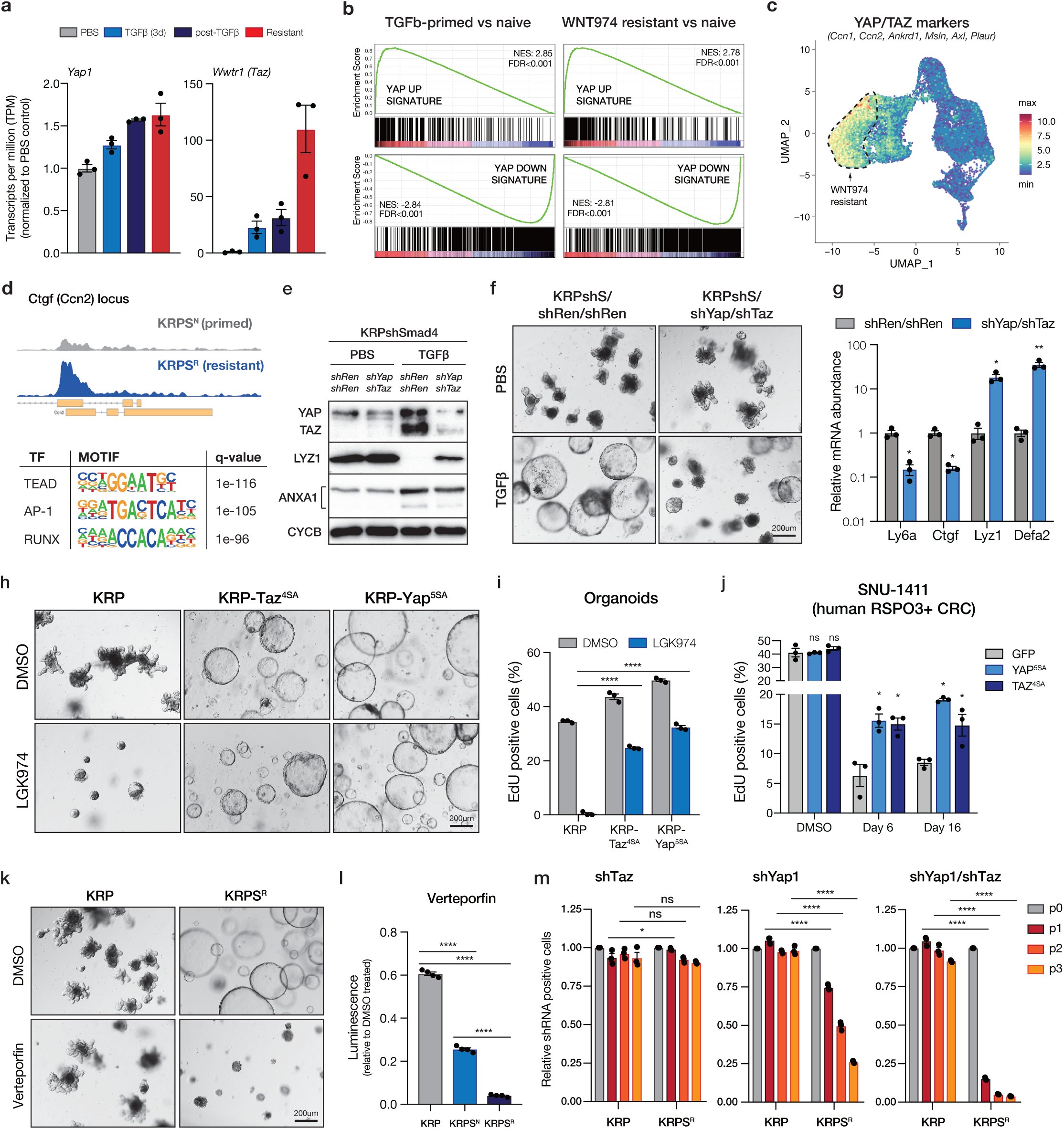
Yap/Taz is required for lineage reversion and WNT independence. **a.** Normalized TPM shows upregulation of Yap1 and Taz (Wwtr1) transcripts following TGFb-priming and development of WNT-independence. **b**. GSEA plots showing YAP signature is enriched in post-TGFβ KRPshS cells, compared to PBS-treated isogenic organoids. **c.** UMAP plot of scRNAseq data showing relative mean gene expression of known YAP/TAZ transcriptional targets, highlighting increased YAP activity in WNT974 resistant population. **d**. ATACseq profile showing increased accessibility of Ctgf (Ccn2) locus in WNT974-resistant KRPS compared to WNT974-naive KRPS and HOMER motif analysis from ATACseq showing top 3 known transcription factor binding sites enriched in WNT974-resistant KRPS organoids compared to WNT974-naive cells. **e**. Western blot of whole cell lysates from KRPshS/shRen/shRen and KRPshS/shYap/shTaz organoids treated with PBS or 5ng/mL TGFβ as indicated. **f**. Bright field images of KRPshS/shRen/shRen and KRPshS/shYap/shTaz organoids treated with PBS or 5ng/mL TGFβ as indicated. **g**. qRT-PCR analysis of gene expression in KRPshS/shRen/shRen and KRPshS/shYap/shTaz organoids treated with PBS or 5ng/mL TGFβ as indicated. **h**. Bright field images of KRP, KRP-TAZS4A and KRP-YAPS5A organoids treated with DMSO or 500nM WNT974 for 4 days. **i**. EdU flow cytometry on KRP, KRP-TAZS4A and KRP-YAPS5A organoids treated with DMSO or 500nM WNT974 for 4 days. **j**. EdU flow cytometry quantitation on SNU1411 human RSPO3 fusion CRC cells expressing control vector (GFP) or YAPS5A or TAZS4A treated with WNT974 in 3D Matrigel culture, as indicated. Data represent treatment of 3 independent transductions (n=3, *p<0.05, two-way ANOVA, with Tukey’s correction). **k**. Bright field images of KRP and KRPS WNT974-resistant line treated with 1uM Verteporfin for 48h. **l**. Cell viability assay on KRP, KRPS WNT974-naive line and KRPS WNT974-resistant line treated with 1uM Verteporfin for 48h. **m.** Cell competition assays on KRP and KRPS WNT974-resistant line transduced with shTaz, shYap1 and shYap1/shTaz.

Together, these molecular data suggest that the transcriptional reprogramming was likely driven by direct YAP/TAZ/TEAD DNA-binding and that robust YAP/TAZ activity may be critical for the establishment of WNT independence. YAP/TAZ induction was also clearly apparent in vivo, with KRPS tumors showing markedly elevated nuclear YAP/TAZ staining, compared to KRP and KRPT genotypes (Supplementary Figure 9c). We further observed prominent nuclear active (non-phosphorylated) YAP in a subset (5/8) of human RSPO2/RSPO3 fusion CRCs, all of which contained concomitant mutations in KRAS (or BRAF) and TP53 (Supplementary Figure 10).

To directly assess whether YAP/TAZ signaling was the key driver of WNT independence downstream of TGFβ, we silenced YAP and TAZ in TGFβ-naïve KRPshS organoids using tandem shRNAs. YAP/TAZ silencing in these cells had no obvious impact on morphology or organoid growth under basal conditions, but like SMAD2/3 knockdown, disrupted the transition to spheroid morphology, and blocked key molecular and transcriptional changes associated with lineage reversion (Figure 6e-g). YAP/TAZ activity was not only required for spheroid formation and transcriptional changes downstream of TGFβ, but was sufficient to induce WNT independence. Expression of either stabilized YAP (YAP^S5A^) or TAZ (TAZ^S4A^) in WNT-dependent KRP organoids was sufficient to downregulate adult intestinal lineage markers, upregulate fetal intestinal signature genes and drive spheroid transformation in the absence of TGFβ priming (Figure 6h, Supplementary Figure 11). Moreover, KRP-YAP^S5A^ and KRP-TAZ^S4A^ organoids were refractory to WNT inhibition with WNT974, showing only a moderate decrease in EdU incorporation (Figure 6i), comparable to KRPS^R^ organoids (Figure 2b). Similarly, expression of YAP^S5A^ or TAZ^S4A^ in the RSPO3 fusion-positive human CRC cell line SNU1411, led to a 2.5-fold increase in proliferation following treatment with WNT974 (Figure 6j).

To determine whether YAP/TAZ-mediated signaling was required for maintenance of WNT-independence, we treated KRP and KRPS^R^ cells with Verteporfin (VP), which disrupts the YAP/TAZ-TEAD protein interaction (37), in addition to other cellular targets. Remarkably, while VP had no effect on the morphology and caused a modest decrease in growth of WNT-dependent KRP organoids, it was profoundly toxic to KRPS^R^ cells, significantly reducing cell viability and completely blocking spheroid growth (Figure 6k-l). To ensure that this effect was due to the disruption of YAP signaling and not another effect of VP treatment, we silenced YAP, TAZ, or both YAP and TAZ in KRP and KRPS^R^ cells and measured proliferation by fluorescence-based competition assay. TAZ knockdown alone had very little impact on the proliferation of organoids relative to untransduced cells, while YAP silencing had a more dramatic effect on KRPS^R^ cells (Figure 6m). Consistent with the effect of VP treatment, combined YAP/TAZ suppression induced a rapid and near-complete depletion of shRNA-expressing cells by 6 days (P2) (Figure 6m). In contrast, YAP/TAZ knockdown had no effect on the proliferation of KRP cells (Figure 6m), highlighting the switch from WNT to YAP dependency following TGFβ-primed transcriptional rewiring and WNT independence.

Together, these data suggest that YAP/TAZ transcriptional activity is the central requirement for TGFβ-primed lineage reversion, and that following the transition to WNT independence, YAP/TAZ signaling becomes essential for survival and proliferation.

## DISCUSSION

WNT pathway hyperactivation is the most frequent initiating in human colorectal cancer and thought to be one of the most important molecular drivers in this disease. Here, we show that intestinal tumor cells carrying common cancer-associated genetic alterations can become primed to evade targeted WNT inhibition and rapidly evolve to become completely WNT independent. WNT independence is initiated by canonical TGFβ signaling, which drives YAP/TAZ-dependent transcriptional reprogramming and lineage reversion. Consequently, WNT independent cells become reliant on YAP/TAZ signaling and can be selectively depleted by genetic or pharmacologic targeting of YAP.

The role of YAP in intestinal biology is complex. YAP is required for early phases of tumor initiation in APC mutant cells (38,39), but in normal intestinal epithelium, YAP induction leads to crypt loss and intestinal dysfunction (40). Likewise, activating genetic alterations in YAP, TAZ and TEAD are rare in colorectal cancer, although YAP signaling can be important for proliferation and metastasis in WNT-driven CRC xenografts (41). Our data show that YAP/TAZ signaling can be potent driver of survival and proliferation in intestinal cancer cells, independent of WNT activation. This mirrors a recently described role for YAP/TAZ in normal tissue regeneration following colitis (33) and/or irradiation-induced mucosal damage (31). In both cases, YAP-mediated repair is associated with upregulation of fetal markers, though it is unclear whether the regenerating epithelium, like fetal organoids (28), are WNT-independent. Taken together, it seems likely that TGFβ-priming in KRPS/shAKPS cells promotes WNT-independence by hijacking a normal physiological wound-repair process to drive lineage reversion.

Despite the similarities, resistance to WNT blockade in cancer cells has two critical and fundamental differences to the wound-repair program. First, TGFβ-priming, unlike injury-induced YAP activation, is dependent on the presence of multiple oncogenic mutations (e.g. KRAS/BRAF, p53 and SMAD4), indicating that this is mode of reprogramming is likely cancer-specific. Second, YAP-mediated wound repair is a tightly controlled and transient process (31,33), while lineage reversion of KRPS/AKPS cells is irreversible. Even following months of culture in the absence of WNT suppression, WNT-independent organoid cultures remain resistant to WNT974 and reliant on YAP/TAZ for survival. Understanding the factors that dictate the differences between reversible YAP activation in injury repair and permanent YAP-reprogramming in WNT inhibitor resistance may identify key cellular switches that could be harnessed to prevent or reverse WNT-independence in therapy refractory tumors. For instance, while blocking the TGFβ receptor or restoring SMAD4 has no impact on the maintenance of WNT-independence, it is possible that cytokine-dependent YAP induction guides reprogramming on a different course to transient wound repair. Further, disruption of p53 has been shown to impact cellular reprogramming and lineage plasticity in multiple settings (42-45), but it is not clear exactly what role p53 loss plays in the initiation or maintenance of fetal lineage reversion in the gut. Interestingly, p53 disruption is extremely common in patients with colitis-associated CRC, whose tumors show relatively infrequent WNT activating mutations (46). It is tempting to speculate that induction of an inflammation-induced YAP regenerative program in p53 mutant cells would enable WNT-independent cancer growth. It is notable that TP53 (and KRAS or BRAF) mutations were present in each human RSPO3 fusion CRC case we identified with nuclear (active) YAP1. Interestingly, nuclear YAP1 did not correlate precisely with SMAD4 status, implying that SMAD4 loss is not an absolute requirement for YAP/TAZ activation in CRC, which can be induced by factors other than TGFβ (41). Hence, while we describe one mechanism of lineage reversion and WNT-independence, we suspect that there will be other genetic combinations, contexts, and/or extracellular stimuli that induce a similar WNT-independent pro-tumorigenic outcome. Previous studies have identified elevated expression of a variety of YAP and fetal-like markers in human CRC, including ANXA1 (47), SPP1 (48), and MSLN (49,50). Whether these markers alone can distinguish WNT independent or ‘primed’ human tumors is unknown.

The work described here was guided by the clinical genetics of CRC, and in particular, RSPO fusion CRC. Due to the paucity of pre- and post-treatment clinical samples, we exploited engineered murine organoids to interrogate the genetics of WNT dependence and WNT-targeted therapy response, but this work has obvious implications for clinical application of WNT therapies. To date, WNT-targeted drugs have performed poorly in early phase clinical studies, owing to on-target dose-limiting toxicity, including bone fractures (51). Recent work has shown that combination of RANKL and PORCN inhibitors provide potent WNT pathway suppression while avoiding adverse bone-related effects (23). Based on this data, new combination trials have been launched, specifically targeting RSPO3 fusion CRC. It is too early to know the outcome of these new trials, but given the prevalence of p53, SMAD4, and RAS/RAF alterations in RSPO fusion cancers, it will be critical to pay close attention to how genotype influences clinical tumor behavior. Ultimately, such genetic biomarkers may serve to further stratify patients to improve clinical outcomes.

## Acknowledgements

We thank Wouter Karthaus, Charles Sawyers, and Darjus Tschaharganeh for technical and experimental advice. We thank Francisco Barriga, Michal Nagiec, Ashley Laughney, and J. Joshua Smith for advice and comments on preparation of the manuscript. This work was supported by a Research Scholar Award from the American Cancer Society (RSG-17-202-01-TBG), project grant from the NIH/NCI (1R01CA222517-01A1) and a Stand Up to Cancer Colorectal Cancer Dream Team Translational Research Grant (SU2C-AACR-DT22-17). Stand Up to Cancer is a program of the Entertainment Industry Foundation. Research grants are administered by the American Association for Cancer Research, a scientific partner of SU2C. This work was also supported by Starr Cancer Consortium grant I11-0040. We thank the Weill Cornell Genomics Resource Core Facility who performed library preparation and sequencing for WES, RNAseq, and scRNAseq, and the Weill Cornell Epigenomics Core Facility who performed library preparation and ATACseq. MPZ is supported in part by National Cancer Institute (NCI) Grant NIH T32 CA203702. The content is solely the responsibility of the authors and does not necessarily represent the official views of the NIH.

## Author Contributions

TH designed and performed experiments, analyzed data and wrote the paper. SG, FT, MPZ, CM, ZC, JTP, JFH, and RY performed experiments and/ or analyzed data. EK and OE supervised experiments and/or analysis. LED designed and supervised experiments, analyzed data, and wrote the paper.

## METHODS

### Cloning

Sequences encoding guide RNAs (Supplementary Table 2) were cloned into BsmBI site of Lenti-Cas9-Cre (LCC) vector. For tandem guide RNA cloning, U6-sgRNA cassette was amplified from px458 vector, then was ligated into EcoRI site of LCC vector. For shRNA cloning, shRNA (Supplementary Table 2) was cloned into XhoI/EcoRI site of SGEN vector. For tandem shRNA cloning, shRNA was cloned into EcoRI site downstream of the existing shRNA in SGEN vector. For cDNA cloning, cDNA was ligated or Gibson assembled into BglII/MluI site of LMN vector.

### Animal studies

All studies involving animals were approved by the Institutional Animal Care and Use Committee (IACUC) at Weill Cornell Medicine (NY), under protocol number 2014-0038. For Apc restoration studies, animals were fed with Doxycycline chew (200mg/kg) (Harlan Teklad) 1 day before transplantation. Organoids (∼100,000 cells for each flank) were implanted subcutaneously into athymic nude mice. Animals were distributed into “ON Dox” group and “OFF DOX” group when tumors reached a volume of approximately 100mm3. All animals were humanely euthanized after 5 weeks. Tumor samples were weighed and sent for histology. For WNT974 treatment studies, organoids (∼100,000 cells for each flank) were implanted subcutaneously into athymic nude mice. After 2 weeks, animals were randomized for WNT974 treatment. WNT974 (5mg/kg, Selleckchem #S7143) was mixed with 0.5% methylcellulose and 0.1% Tween 80 and then administrated by daily oral gavage for 2 weeks. After treatment, all animals were humanely euthanized. Tumor samples were weighed and sent for histology.

### Immunohistochemistry and immunofluorescence

Tissue, fixed in freshly prepared 4% paraformaldehyde for 24 hours, was embedded in paraffin and sectioned by IDEXX RADIL (Columbia, MO). Sections were rehydrated and unmasked (antigen retrieval) by either: (i) Heat treatment for 5 mins in a pressure cooker in 10mM Tris / 1mM EDTA buffer (pH 9) containing 0.05% Tween 20 or (ii) Proteinase K treatment (200ug/ml) for 10 mins at 37ºC (Lysozyme staining). For immunohistochemistry, sections were treated with 3% H2O2 for 10 mins and blocked in TBS / 0.1% Triton X-100 containing 1% BSA. For immunofluorescence, sections were not treated with peroxidase. Primary antibodies, incubated at 4C overnight in blocking buffer, were: rabbit anti-ki67 (1:100, Sp6 clone, Abcam #ab16667), rabbit anti-KRT20 (1:200, Cell Signaling Technologies, #13063), rabbit anti-Lysozyme (1:1000, Dako, #EC 3.2.1.17), rat anti-BrdU (1:200, Abcam #ab6326), rabbit anti-YAP/TAZ (1:200, Cell Signaling Technologies, #8418), mouse anti-Spp1 (1:200, Santa Cruz Biotechnology, #sc-21742). For immunohistochemistry, sections were incubated with anti-rabbit or anti-rat ImmPRESS HRP-conjugated secondary antibodies (Vector Laboratories, #MP7401) and chromagen development performed using ImmPact DAB (Vector Laboratories, #SK4105). Stained slides were counterstained with Harris’ hematoxylin. For immunofluorescent stains, secondary antibodies were applied in TBS for 1 hour at room temp in the dark, washed twice with TBS, counterstained for 5 mins with DAPI and mounted in ProLong Gold (Life Technologies, #P36930). Secondary antibodies used were: anti-rabbit 568 (1:500, Molecular Probes, #a11036). Images of fluorescent and IHC stained sections were acquired on a Zeiss Axioscope Imager (chromogenic stains), Nikon Eclipse T1 microscope (IF stains). Raw .tif files were processed using FIJI (Image J) and/or Photoshop (Adobe Systems Inc., San Jose, CA) to create stacks, adjust levels and/or apply false coloring.

### Isolation and culture of intestinal organoids

Isolation, maintenance and staining of mouse intestinal organoids has been described previously (52,53). Briefly, for isolation, 15 cm of the proximal small intestine was removed and flushed with cold PBS. The intestine was then cut into 5 mm pieces, vigorously resuspended in 5mM EDTA-PBS using a 10ml pipette, and placed at 4ºC on a benchtop roller for 10 minutes. This was then repeated for a second time for 30 minutes. After repeated mechanical disruption by pipette, released crypts were mixed with 10ml DMEM Basal Media (Advanced DMEM F/12 containing Pen/Strep, Glutamine, 1mM N-Acetylcysteine (Sigma Aldrich A9165-SG)) containing 10 U/ml DNAse I (Roche, 04716728001), and filtered sequentially through 100μm and 70μm filters. 1ml FBS (final 5%) was added to the filtrate and spun at 125g for 4 minutes. The purified crypts were resuspended in basal media and mixed 1:10 with Growth Factor Reduced Matrigel (BD, 3542l of the resuspension was plated per well in a 48 well plate and placed in a 37ºC incubator to polymerize for 10 minutes. 250μl of small intestinal organoid growth media (Basal Media containing 50 ng/mL EGF (Invitrogen PMG8043), 100ng/ml Noggin (Peprotech 250-38), and 500 ng/mL R-spondin (R&D Systems, 3474-RS-050, or from conditioned media) was then laid on top of the Matrigel. Where indicated, dox was added to experiments at 500 ng/ml.

For sub-culture and maintenance, media was changed on organoids every two days and they were passaged 1:4 every 5-7 days. To passage, the growth media was removed and the Matrigel was resuspended in cold PBS and transferred to a 15ml falcon tube. The organoids were mechanically disassociated using a p1000 or a p200 pipette and pipetting 50-100 times. 7 ml of cold PBS was added to the tube and pipetted 20 times to fully wash the cells. The cells were then centrifuged at 125g for 5 minutes and the supernatant was aspirated. They were then resuspended in GFR Matrigel and replated as above. For freezing, after spinning the cells were resuspended in Basal Media containing 10% FBS and 10% DMSO and stored in liquid nitrogen indefinitely.

### Organoid Imaging

For fixed staining, organoids were grown in 40μl of Matrigel plated into an 8-well chamber slide (Lab-Tek II, 154534). Where indicated, 10μM EdU was added to the growth media for 6 hours before fixing. The growth media was removed and the cells were fixed in 4% PFA-PME (50mM PIPES, 2.5 mM MgCl2, 5mM EDTA) for 20 minutes. They were then permeabilized in .5% Triton for 20 minutes and blocked in IF Buffer (PBS, .2% Triton, .05% Tween, 1% BSA) for 1 hour or immediately processed for EdU staining performed according to directions provided with the Click-iT Edu Alexa Fluor 647 Imaging Kit (Invitrogen C10340). For alkaline phosphatase staining, fixed cells were washed twice with TBS and then incubated with the BCIP/NBT Substrate Kit (Vector Laboratories, SK-5400) for 15 minutes in the dark. The chambers were then washed twice with TBS and then imaged using bright field microscopy. For immunofluorescent staining, cells were incubated in primary antibodies overnight in IF buffer: rabbit anti-KRT20 (1:200, Cell Signaling Technologies, #13063), rabbit anti-Lysozyme (1:200, Dako, #EC 3.2.1.17). They were then washed 3 times with TBS .1% Tween. Secondary antibodies (1:1000, same reagents as above) were incubated with or without Alexa-647 Phalloidin (Molecular Probes, #A22287) for 1 hour. The solution was removed and DAPI in PBS was added for 5 minutes and washed twice with TBS .1% Tween. The chambers were then removed and cover slips were mounted using Prolong Gold antifade medium (Invitrogen P36930). Images were acquired using Zeiss LSM 880 laser scanning confocal microscope, and Zeiss image acquiring and processing software. Images were processed using FIJI (Image J) and Photoshop CS5 software (Adobe Systems Inc., San Jose, CA).

### Organoid transfection

Murine small intestinal organoids were cultured in transfection medium containing CHIR99021 (5μM) and Y-27632 (10μM for 2 days prior to transfection. Single cells suspensions were produced by dissociating organoids with TrypLETM express (Invitrogen #12604) for 5 min at 37°C. After trypsinization, cell clusters in 300μl transfection medium were combined with 100μl DMEM/F12-lipofectamine2000 (Invitrogen #11668)-DNA mixture (97ul-2ul-1ug), and transferred into a 48-well culture plate. The plate was centrifuged at 600g at 32°C for 60 min, followed by another 6h incubation at 37°C. The cell clusters were spun down and plated in Matrigel. For selecting organoids with Ptprk-Rspo3 rearrangements, exogenous RSPO1 was withdrawn 1 week after transfection. Organoids were cultured in medium containing Nutlin3 (5μM) and TGFβ1 (5ng/mL) for 1 week to select for p53 loss and Smad4 loss.

### Protein analysis

Small intestine organoids were grown in 300μl of Matrigel in 1 well of 6-well plate for 4 days post-passage. Organoids were then recovered from the Matrigel using cell recovery solution (Corning #354253). Organoid pellets were lysed with RIPA buffer. Antibodies used for Western blot were: mouse anti-p53 (1:1000, Cell Signaling Technologies, #2524), rabbit anti-Smad4 (1:1000, Cell Signaling Technologies, #46535), rabbit anti-Smad2/3 (1:1000, Cell Signaling Technologies, #8685), rabbit anti-YAP (1:1000, Cell Signaling Technologies, #14074), rabbit anti-YAP/TAZ (1:1000, Cell Signaling Technologies, #8418), rabbit anti-Runx2 (1:1000, Cell Signaling Technologies, #12556), mouse anti-E-Cadherin (1:1000, Cell Signaling Technologies, #14472), rabbit anti-phospho-Akt (Ser473) (1:1000, Cell Signaling Technologies, #4060), rabbit anti-N-Myc (1:1000, Cell Signaling Technologies, #84406), rabbit anti-c-Jun (1:1000, Cell Signaling Technologies, #9165), rabbit anti-Junb (1:1000, Cell Signaling Technologies, #3753), rabbit anti-ETS-1 (1:1000, Cell Signaling Technologies, #14069), rabbit anti-GAPDH (1:1000, Cell Signaling Technologies, #5174), rabbit anti-Cyclophilin B (1:1000, Cell Signaling Technologies, #43603), rabbit anti-pan-TEAD (1:1000, Cell Signaling Technologies, #13295), rabbit anti-phospho-Smad2 (Ser465/467) (1:1000, Cell Signaling Technologies, #3108), mouse anti-phospho-SAPK/JNK (Thr183/Tyr185) (1:1000, Cell Signaling Technologies, #9255), rabbit anti-phospho-p38 MAPK (Thr180/Tyr182) (1:1000, Cell Signaling Technologies, #4511), mouse anti-TEAD4 (1:1000, Abcam, #ab58310), rabbit anti-phospho-Smad3 (Ser423/425) (1:1000, Abcam, #ab52903).

### Flow Cytometry

Organoids were pulsed with 10uM EdU for 3 hours in regular tissue culture incubator. Cells were pelleted and trypsinized using 0.25% Trypsin-EDTA (ThermoFisher scientific #25200056) at 37 degree for 5 minute, re-suspended using FACS buffer (PBS+2%FBS) then filtered through cell strainer (Corning #352235). EdU was stained using Click-iTTM EdU Alexa FluorTM 647 flow cytometry assay kit (C104424) following the manufacturer instructions. Flow cytometry was conducted on Invitrogen Attune NxT following the manufacturer instructions.

### RNA isolation, cDNA synthesis and QPCR

RNA was extracted using TRIzol according to the manufacturer’s instructions and contaminating DNA was removed by DNase treatment for 10 mins and column purification (Qiagen RNAeasy). cDNA was prepared from 1µg total RNA using qScript reverse transcription kit (Quantabio, #95047). Quantitative PCR detection was performed using SYBR green reagents (Quantabio #101414-288) and specific primers listed in Supplementary Table 2.

### Whole exome sequencing

Each gDNA sample based on Qubit quantification are mechanically fragmented on an Covaris E220 focused ultrasonicator (Covaris, Woburn, MA, USA). Two hundred ng of sheared gDNA were used to perform end repair, A-tailing and adapter ligation with Agilent SureSelect XT (Agilent Technologies, Santa Clara, CA) library preparation kit following the manufacturer instructions. Then, the libraries are captured using Agilent SureSelectXT Mouse All Exon probes, and amplified. The quality and quantities of the final libraries were checked by Agilent 2100 Bioanalyzer and Invitrogen Qubit 4.0 Fluorometer (Thermo Fisher, Waltham, MA), libraries are pooled at 8 samples per lane and sequenced on an Illumina HiSeq 4000 sequencer (Illumina Inc, San Diego, CA) at PE 2×100 cycles. Copy number alterations were identified and plotted using cnvkit (v0.9.6) and single nucleotide varaiant called using MuTect2.

### RNA sequencing

Total RNA was isolated using Trizol, DNAse treated and purified using the RNeasy mini kit (Qiagen, Hilden, Germany). Following RNA isolation, total RNA integrity was checked using a 2100 Bioanalyzer (Agilent Technologies, Santa Clara, CA). RNA concentrations were measured using the NanoDrop system (Thermo Fisher Scientific, Inc., Waltham, MA). Preparation of RNA sample library and RNAseq were performed by the Genomics Core Laboratory at Weill Cornell Medicine. Messenger RNA was prepared using TruSeq Stranded mRNA Sample Library Preparation kit (Illumina, San Diego, CA), according to the manufacturer’s instructions. The normalized cDNA libraries were pooled and sequenced on Illumina NextSeq500 sequencer with single-end 75 cycles.

### RNAseq analysis

The quality of raw FASTQ files were mapped to mouse reference GRCm38 using STAR two-pass alignment (v2.4.1d; default parameters) (54), and transcript abundance estimates were performed using Kallisto (55), aligned to the same (GRCm38) reference genome. Kallisto transcript count data for each sample was concatenated, and transcript per million (TPM) data was reported for each gene after mapping gene symbols to ensemble IDs using R packages, “tximport”, tximportData”, “ensembldb”, and “EnsDb.Mmusculus. v79”. Differential gene expression was estimated using DESeq2 (56). For data visualization and gene ranking, log fold changes were adjusted using lfcShrink in DESeq2, to minimize the effect size of poorly expressed genes. GSEA analysis (v3.0) was performed on pre-ranked gene sets from differential expression between control and treated groups. We used R (v3.6.1) and R Studio (v1.2.1335) to create all visualizations, perform hierarchical clustering and principal component analysis. Volcano plots, heatmaps and other visualizations were produced using the software packages: Enhanced Volcano (https://bioconductor.org/packages/devel/bioc/html/EnhancedVolcano.html), pheatmap (https://cran.r-project.org/web/packages/pheatmap/index.html), ggplot2 (https://cran.r-project.org/web/packages/ggplot2/index.html)

### ATAC sequencing

ATAC-seq library preparation, sequencing and post-processing of the raw data was performed at the Epigenomics Core at Weill Cornell Medicine by using the OMNI-ATAC-seq method previously described (57). Briefly, organoids were dissociated from Matrigel using cell recovery solution (Corning #354253), and 50,000 live cells were spun down and incubated 3 min at 4oC in 25µl of a detergent buffer containing 0.2% Igepal CA-630 (Sigma-Aldrich, St.Louis, MA), 0.2% Tween 20 (Sigma-Aldrich, St.Louis, MA) and 0.02% Digitonin (Promega Corporation, Madison, WI). Nuclei were centrifuged at 500 g for 10 min and immediately resuspended in 25 µl of buffer containing 2.5 µl of Tn5 transposase (Illumina, Inc., San Diego, CA cat # 15027865) for a 30 min incubation at 37oC. Fragments generated by the Tn5 transposase were purified using the DNA Clean and Concentrate kit from Zymo Research (Zymo Research, Irvine, CA cat #D4014). Uniquely indexed libraries were obtained by amplification of the purified fragments with indexed primers using 9 cycles of PCR (5 min × 72 oC, 5 cycles each 10 sec × 98 oC, 30 sec × 63 oC, 1 min × 72 oC). Resulting libraries were subjected to a two-sided size clean up using SPRI beads (Beckman Coulter, Brea, CA) to obtain sizes between 200-1000bp, and pooled for sequencing. The pool was clustered at 9 pM on a pair end read flow cell and sequenced for 50 cycles on an Illumina HiSeq 2500 to obtain ∼40M reads per sample. Primary processing of sequencing images was done using Illumina’s Real Time Analysis software (RTA) as suggested by the Illumina. CASAVA 2.17 software was used to perform image capture, base calling and demultiplexing of the raw reads. FASTQ files generated were aligned to the mouse mm10 genome build using the BWA aligner.

### ATACseq analysis

Raw ATAC-seq reads were processed using ENCODE atac-seq-pipeline(58). This includes Bowtie2(59) alignment against mm10 and MACS2(60) as peak caller. Duplicated reads and reads from mitochondria were filtered out. Peaks for all samples were merged together into a union of peaks, and the ones that existed in at least two samples were kept for further analysis. Peaks were annotated using the ChIPseeker (61) “annotatePeak” function with “TxDb.Mmusculus.UCSC.mm10.knownGene”. Numbers of reads mapped to each peak were counted with featureCounts (62), and raw read counts were normalized using rlog in DESeq2(56). Differentially accessible peaks were defined as peaks with p.adj smaller than 0.05 and absolute value of log2(FC) >1. Known and de novo motifs were idetified using findMotifsGenome.pl from Homer(63) with ‘-size given -mask’ was applied on the differentially accessible peaks, with non-differentially accessible peaks as background.

### Single cell RNA sequencing

Organoids were dissociated from Matrigel using cell recovery solution (Corning #354253), then trypsinized using 10X trypsin (ThermoFisher Scientific #15090046) for 15 minutes at 37C. Single cells were resuspended using organoid culture medium and stained with DAPI. DAPI negative single cells were sorted using BD Aria II FACS machine with 130um nozzle. Sorted single cell suspensions were transferred to the Genomics Core Facility at Weill Cornell Medicine to proceed with the Chromium Single Cell 3’ Reagent Kit v3 (10x Genomics, product code # 1000075) using 10X Genomics’ Chromium Controller. A total of 10,000 cells were loaded into each channel of the Single-Cell A Chip to target 5000 cells in the end. Briefly, according to manufacturer’s instruction, the sorted cells were washed with 1x PBS + 0.04% BSA, counted by Bio-Rad TC20 Cell Counter, and cell viability was assessed and visualized. A total of 10,000 cells and Master Mixes were loaded into each channel of the cartridge to generate the droplets on Chromium Controller. Beads-in-Emulsion (GEMs) were transferred and GEMs-RT was undertaken in droplets by PCR incubation. GEMs were then broken and pooled fractions are recovered. After purification of the first-strand cDNA from the post GEM-RT reaction mixture, barcoded, full-length cDNA was amplified via PCR to generate sufficient mass for library construction. Enzymatic fragmentation and size selection were used to optimize the cDNA amplicon size. TruSeq Read 1 (read 1 primer sequence) was added to the molecules during GEM incubation. P5, P7, a sample index, and TruSeq Read 2 (read 2 primer sequence) were added via End Repair, A-tailing, Adaptor Ligation, and PCR. The final libraries were assessed by Agilent Technology 2100 Bioanalyzer and sequenced on Illumina NovaSeq sequencer with pair-end 100 cycle kit.

### scRNAseq analysis

Reads were pseudoaligned and transcript abundances estimated using kallisto (v0.46.0), and gene count matrices were produced using bustools (v0.39.3) (64) and BUSpaRse. Count matrices were processed and analysed using the Seurat package (v3.1.1) in R (v3.6.1) and R Studio (v1.2.1335). Low quality cells (Less than 1000 identified genes and greater than 10% mitochondrial reads) were removed from the dataset after normalization. Individual scRNAseq datasets were merged and clusters were resolved on the combined data. Expression of cell type markers were plotted using FeaturePlot by calculating the mean expression of the genes indicated for each individual cell.

### Data availability

Raw exomeSeq, RNAseq, ATACseq and scRNAseq data have been deposited in the sequence read archive (SRA) under accession PRJNA578488.

---

**Supplementary Figure 1.**
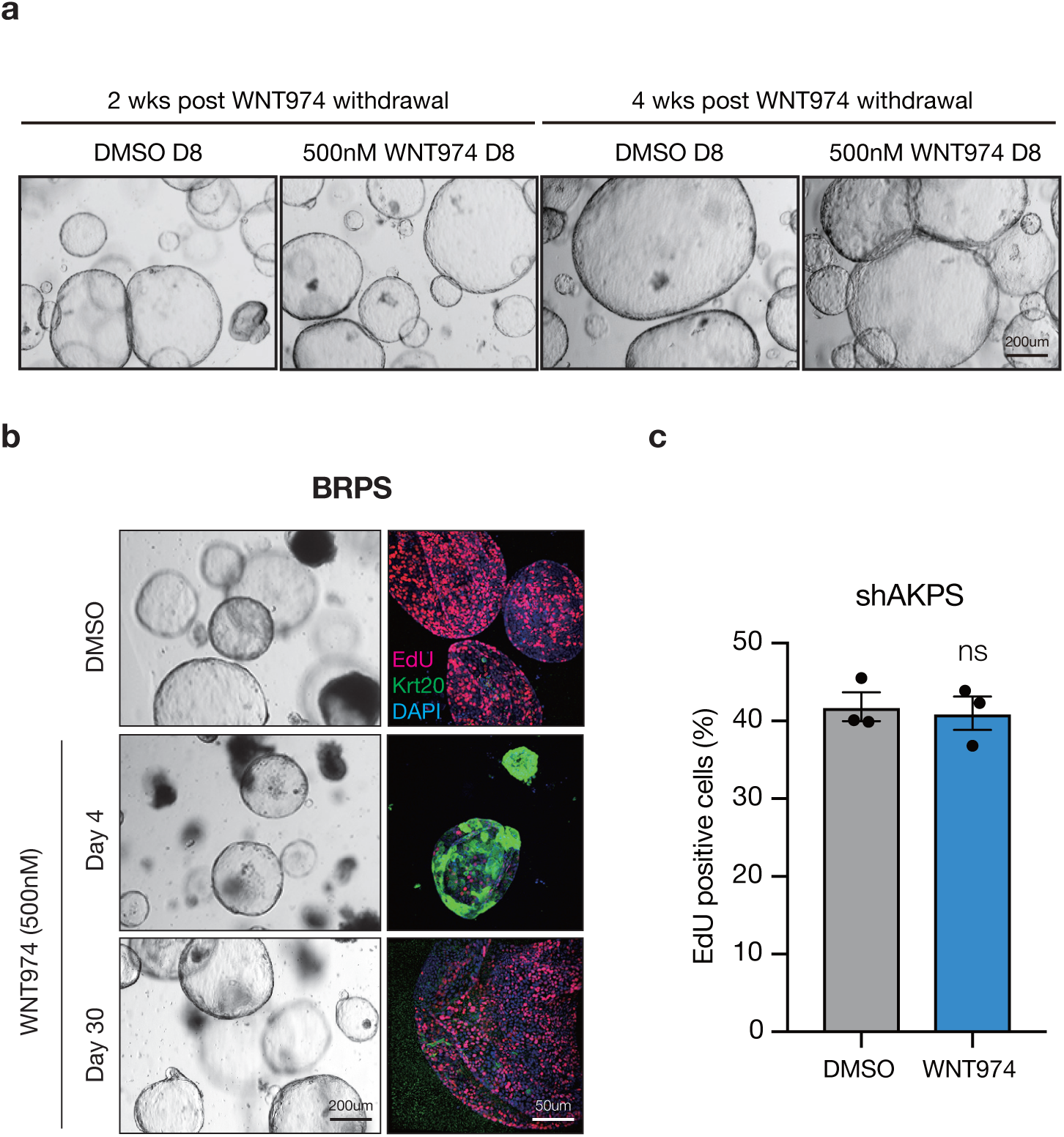
Kras/Braf, p53 and Smad4 mutations enable sustained Wnt independence. **a.** Bright-field images of WNT974-selected out WNT independent KRPS cells show no response when they are re-challenged with WNT974 two weeks and four weeks after WNT974 withdrawal, suggests that these cells are locked in WNT independent state. **b**. Bright field and immunofluorescent images of BRPS organoids treated with DMSO or WNT974. **c**. EdU flow cytometry of Apc restoration resistant shAKPS organoids treated with DMSO or 500nM WNT974 for 4 days. Error bars show +/- SEM, p-value calculated by two-sided t-test with Welsh’s correction.

**Supplementary Figure 2.**
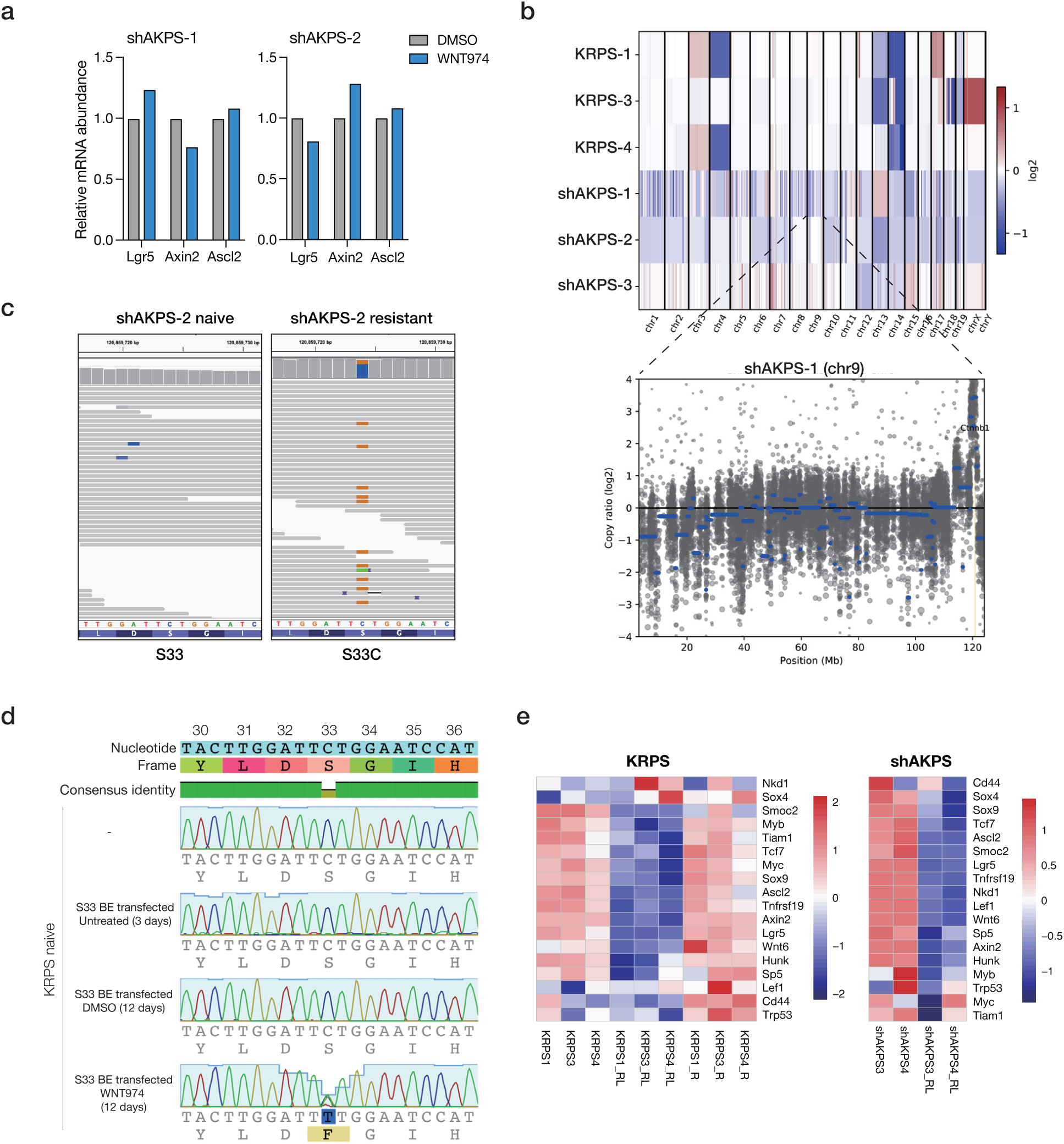
Downstream activation of WNT signaling in two WNT974-resistant organoid lines. **a.** mRNA abundance of WNT target genes in shAKPS1 and shAKP2 organoid lines, measured by qPCR**. b.** Copy number alterations in WNT974-resistant KRPS lines and Apc restoration-resistant shAKPS lines (upper), Ctnnb1 amplification in shAKPS-1 is highlighted (lower). **c**. IGV plot showing hte emergence of a Ctnnb1 hotspot mutation in resistant shAKPS2 population. **d**. De novo generation of an S33F mutation in Ctnnb1 by base editing in KRPS naive organoids, enables outgrowth in the presence of WNT974. **e**. Heatmap of canonical WNT targets from RNAseq data showing WNT signaling is suppressed in WNT974-resistant KRPS organoids on drug (KRPS_RL) compared to drug naive KRPS, but is restored after drug withdrawal (KRPS_R) (left); Similarly, WNT signaling is suppressed in Apc restoration-resistant shAKPS organoids (shAKPS_RL) compared to Apc-depleted shAKPS (right).

**Supplementary Figure 3.**
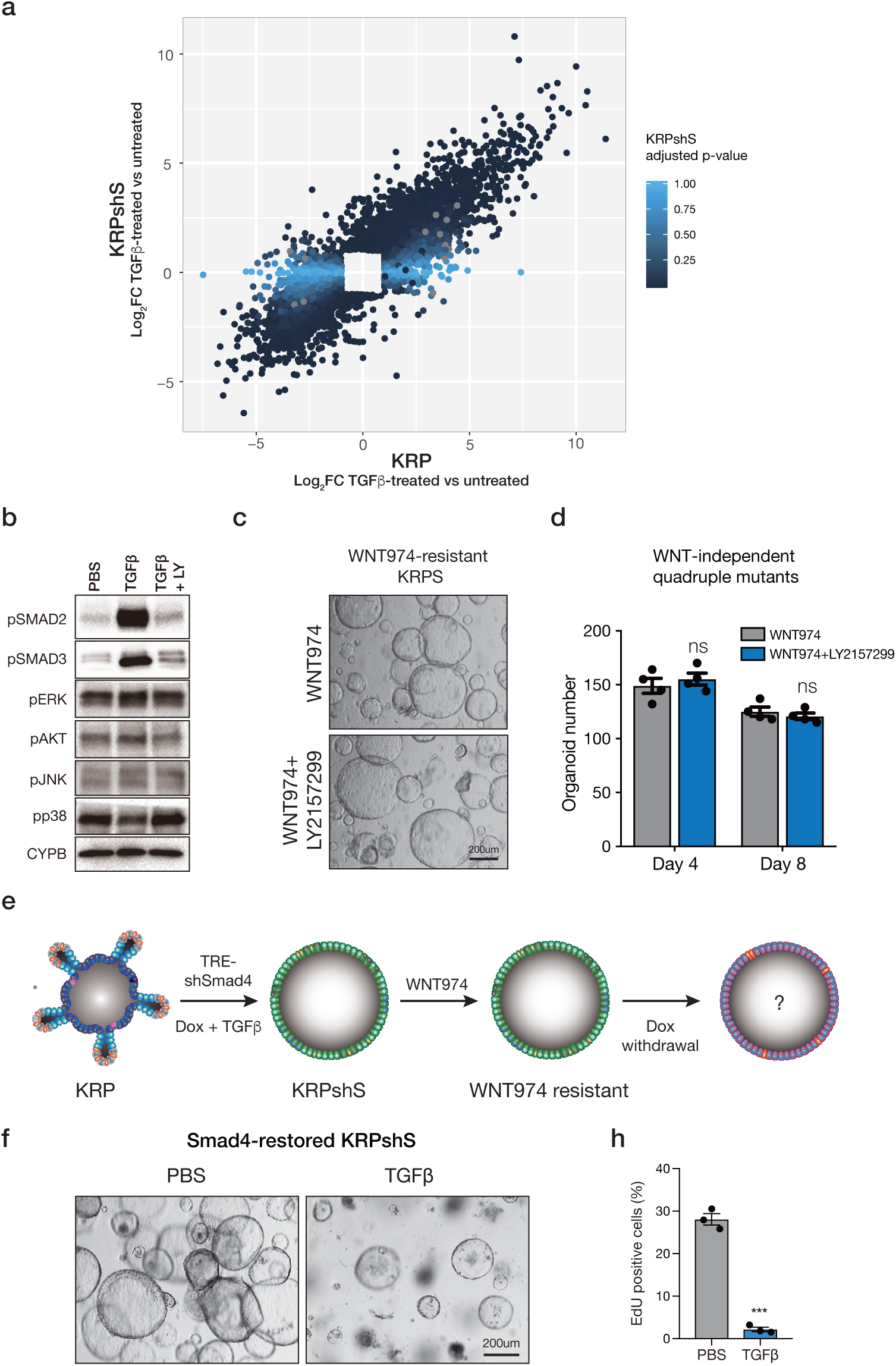
TGFβ or Smad4 loss is not required for the maintenance of WNT independence. **a**. Scatter plot showing Log2 fold change (Log_2_FC) of genes significantly up or downregulated (Log_2_FC >1, padj < 0.01) following 3 days of TGFβ treatment (5ng/ml) in either KRP or KRPshS organoids. Color indicates the adjusted p-value for each given gene in KRPshS cells. Gene expression changes in KRP and KRPshS cells are largely similar, despite the depletion of Smad4 protein. **b.** Western blot of KRPshS organoids treated with PBS, TGFβ, or TGFβ/LY2157299 for 24 hours. **c**. Bright field images of WNT974-resistant KRPS organoids treated with 500nM WNT974 or 500nM WNT974 plus 10uM LY2157299 for 8 days. **d.** Quantification of WNT974-resistant KRPS organoids treated with 500nM WNT974 or 500nM WNT974 plus 10uM LY2157299 for 4 days and 8 days. **e**. Schematic diagram of testing the requirement of Smad4 loss in WNT-independent cells using inducible shSmad4. **f.** Bright field images of Smad4-restored WNT-independent KRPshSmad4 treated with PBS or TGFβ for 4 days. **h.** EdU flow cytometry of Smad4-restored WNT-independent KRPshSmad4 treated with PBS or TGFβ for 4 days. Error bars represent +/- SEM, *p<0.001, two-sided t-test with Welsh’s correction.

**Supplementary Figure 4.**
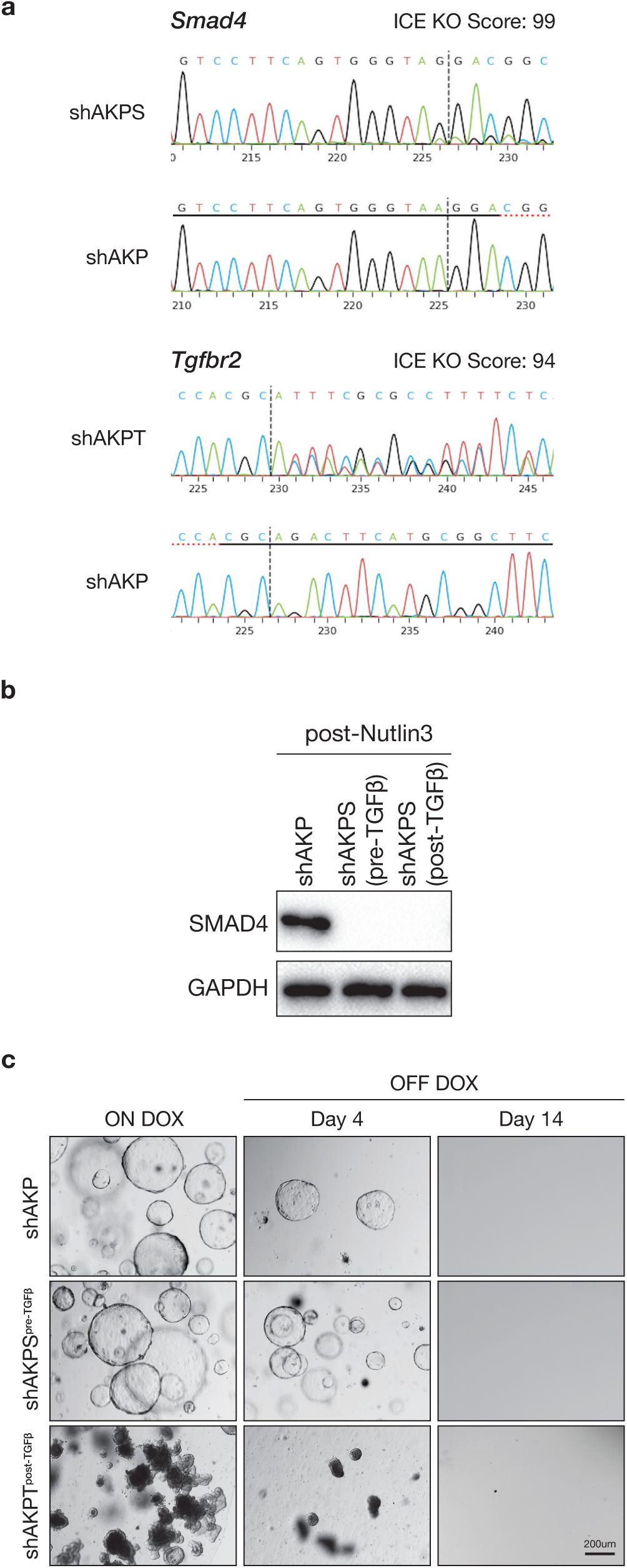
Validation of TGFb-naive shAKPS cells for transplantations. **a.** Synthego ICE sequence analysis of Nutlin3-selected, TGFβ-naive shAKPS and TGFβ-treated shAKPT organoids before transplantation. sgRNA targeting sequence is highlighted by black solid line, PAM sequence is highlighted by red dash line, CRISPR cutting site is labeled by vertical black dash line. **b.** Western blot shows depletion of SMAD4 in both pre- and post-TGFβ organoids. **c.** Bright field images of shAKP, shAKPS (pre-TGFβ) and shAKPT (post-TGFβ) demonstrating that they were unable to escape Apc restoration prior to *in vivo* transplantation.

**Supplementary Figure 5.**
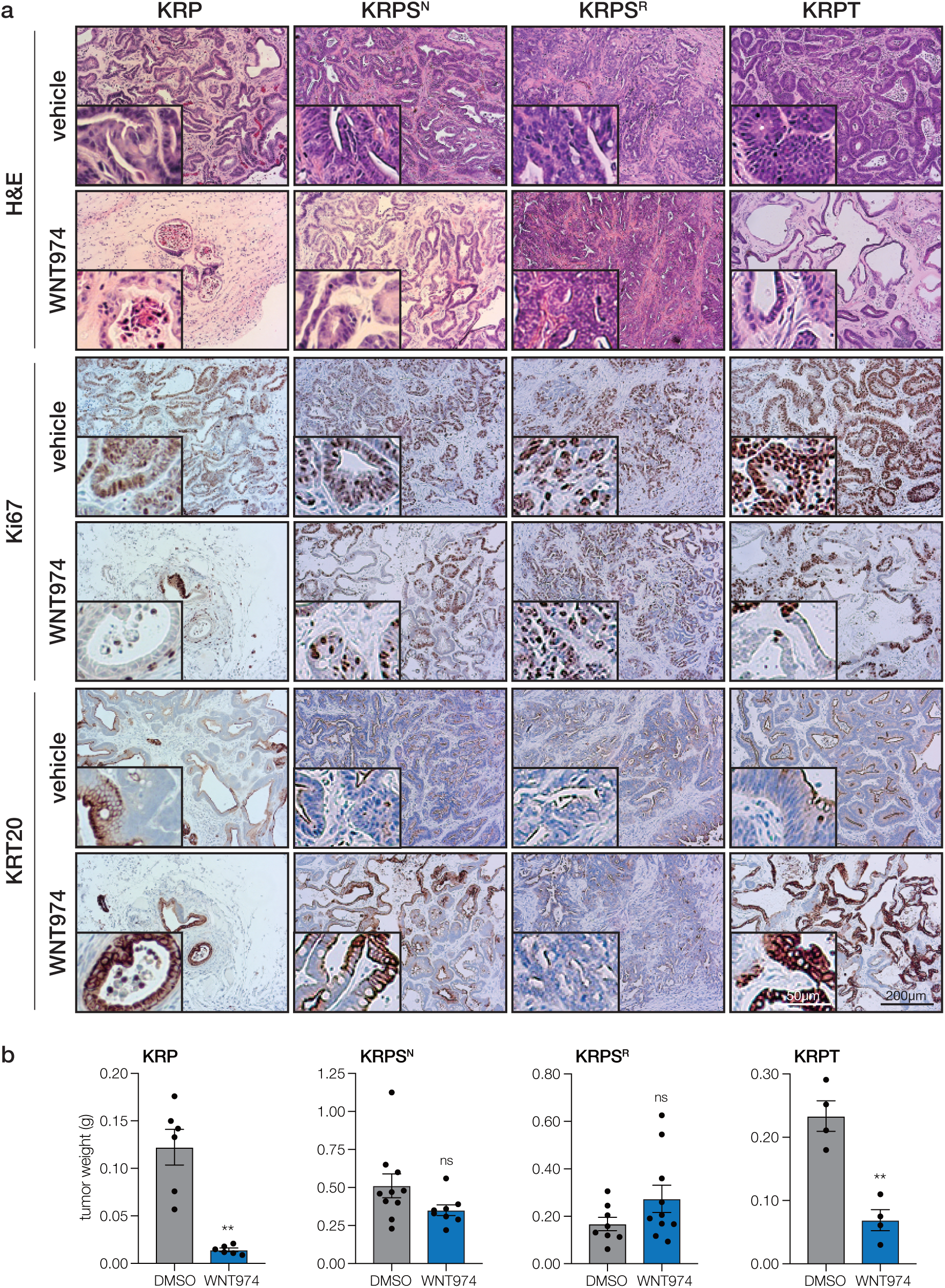
In vivo WNT974 treatment response mirrors in vitro response. a. Immunohistochemical stains of organoid transplants treated with vehicle or WNT974 for 2 weeks. b. Quantification of tumors weight. Error bars represent +/- SEM, n≥4 p-values calculated by two-sided t-test, with Welsh’s correction

**Supplementary Figure 6.**
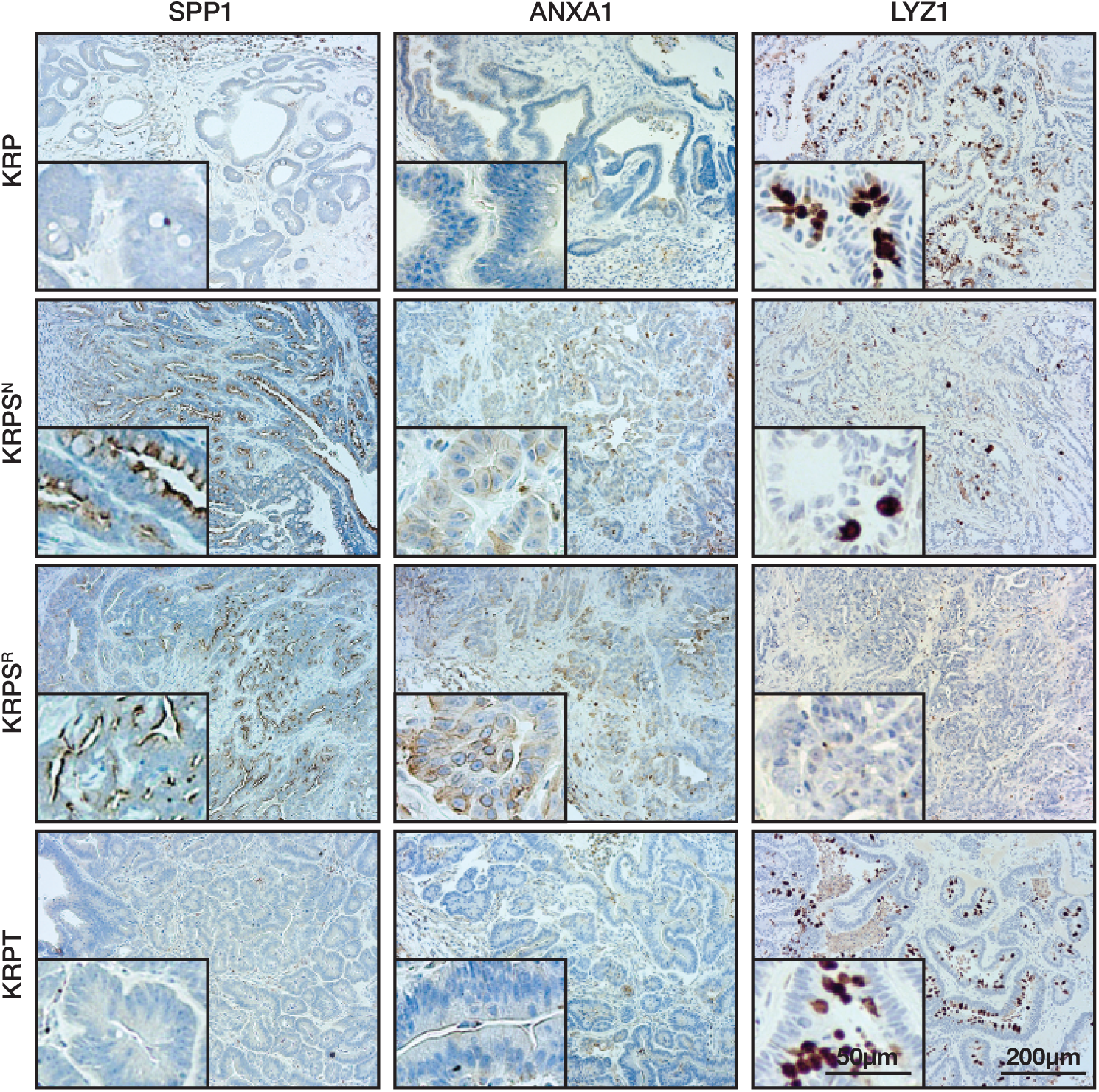
WNT-independent KRPS organoids show fetal-like characteristics *in vivo*. Immunohistochemical stains for Osteopontin (SPP1), Annexin A1 (ANXA1) and Lyzozyme (LYZ1) in organoid-derived tumors, as labeled. TGFβ-primed (KRPS^N^) and WNT974-resistant (KRPS^R^) tumors, but not WNT-dependent KRP and KRPT tumors, show elevated SPP1, ANXA1 stainings and reduced LYZ1-positive Paneth cells.

**Supplementary Figure 7.**
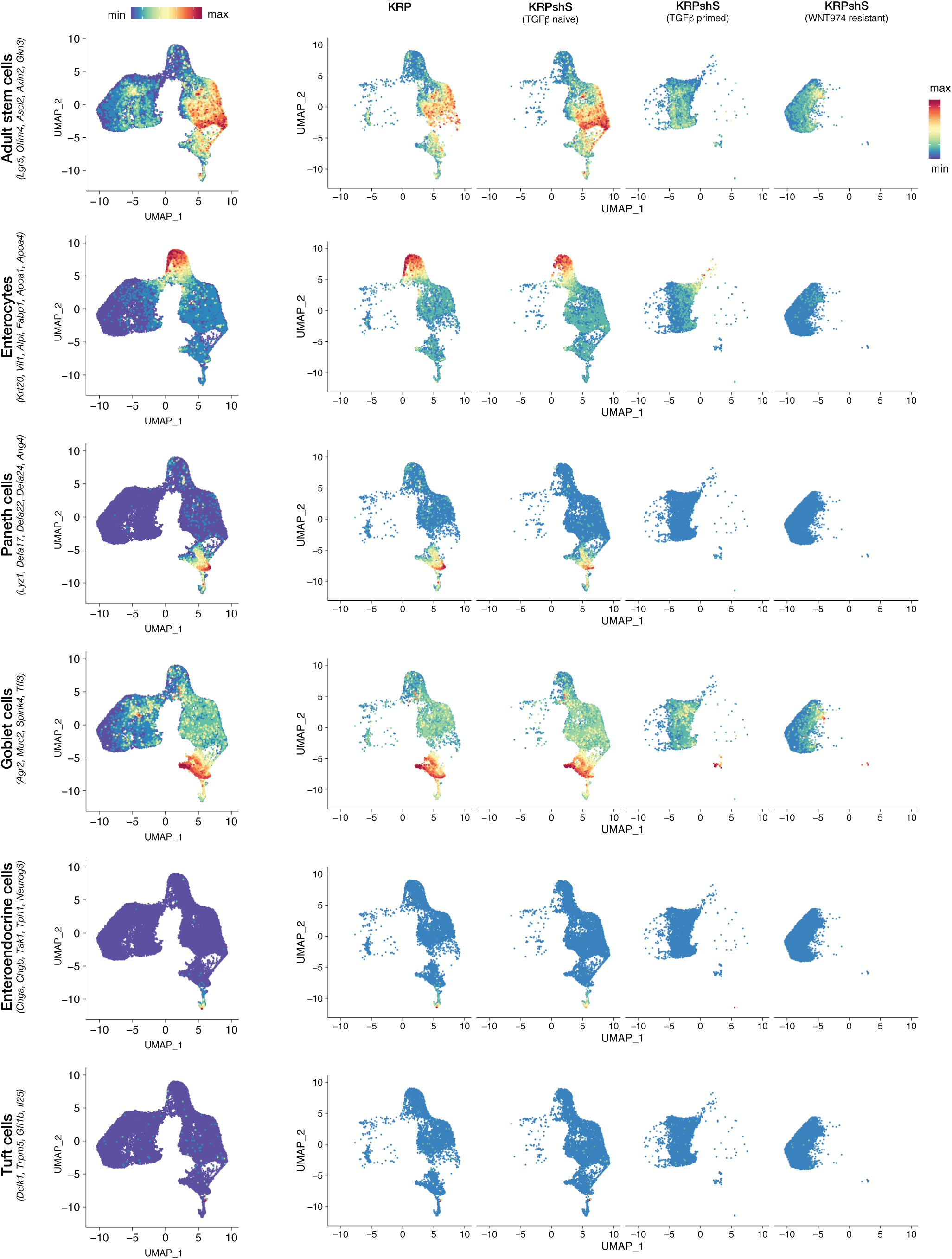
Cluster identity by marker gene expression. Merged (left) and individual UMAP plots showing relative mean expression of marker genes (as labeled). Secretory lineages (goblet and Paneth cells, but not enteroendocrine cells) were grouped as one cluster in our analysis. Tuft cells were not identified as an separate cluster in our analysis, likely owing to the very small number of positive cells.

**Supplementary Figure 8.**
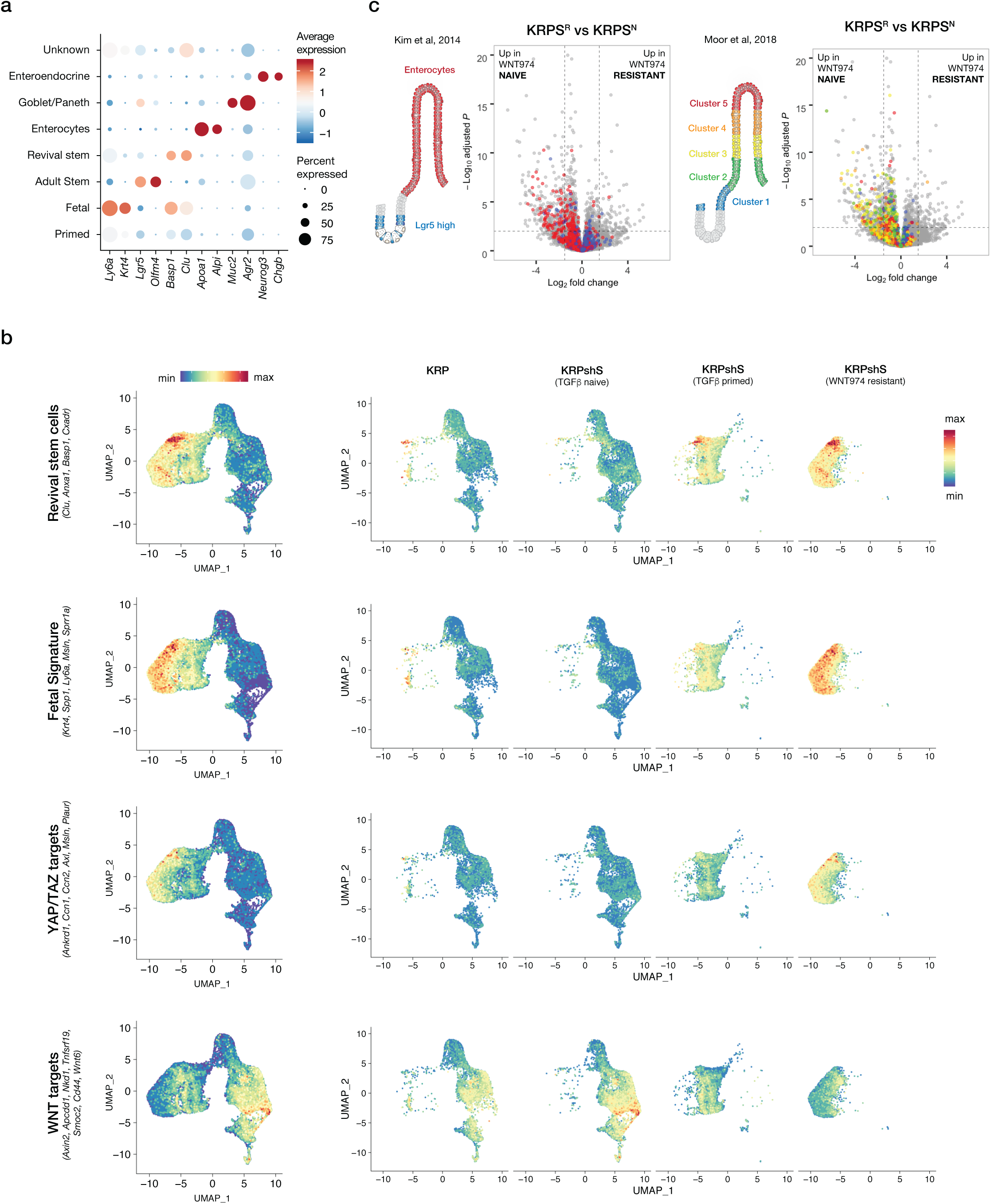
Cluster identity by marker gene expression. **a.** Dot plot showing relative average expression of cell identity markers (x-axis) and the number of cells within each named cluster (y-axis) that express them. **b**. Merged (left) and individual UMAP plots showing relative mean expression of marker genes (as labeled). **c**. Volcano plots of bulk RNAseq of WNT974-naive (KRPS^N^) and WNT974-resistant (KRPS^R^) organoids (all TGFβ-treated), showing reduced expression of enterocyte markers in WNT974-independent organoids.

**Supplementary Figure 9.**
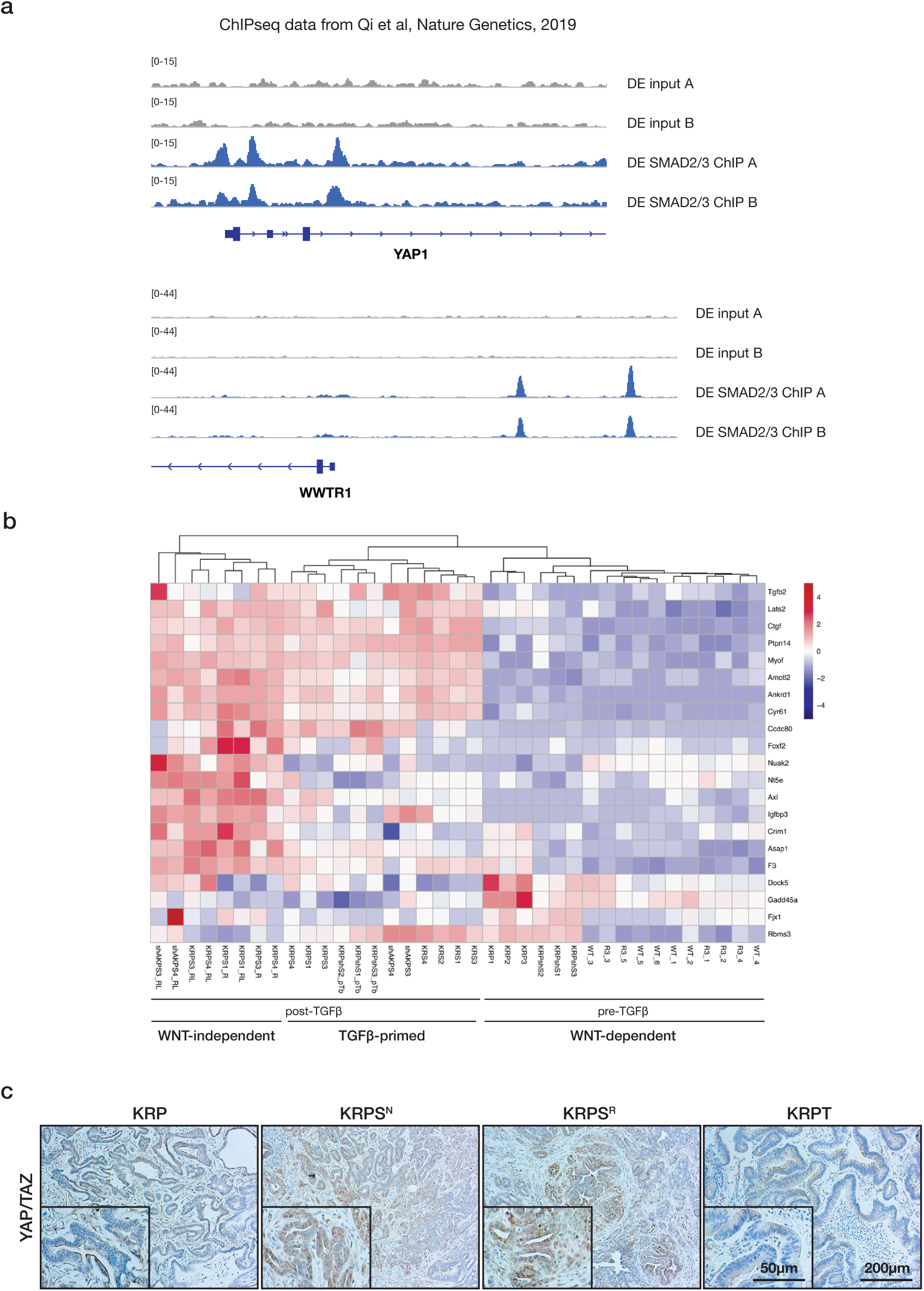
YAP/TAZ transcriptional signature defines TGFβ-primed and WNT independent organoids. **a.** Peak plots of published ChIPseq SMAD2/3 binding on promoter and enhancer regions of *YAP1* and *WWTR1 (TAZ)* in definitive endoderm (DE) derived by *in vitro* hESC differentiation (Qi et al, Nat Gen, 2019). **b.** Unsupervised clustering of organoid lines on YAP/TAZ targets separates all the genotypes into pre-TGFβ, post-TGFβ, and WNT-independent subtypes. **c** *In vivo* tumors from indicated genotypes stained for YAP/TAZ reveal increased and nuclear localization of YAP/TAZ in TGFb-primed (KRPS^N^) and WNT-independent tumors (KRPS^R^).

**Supplementary Figure 10.**
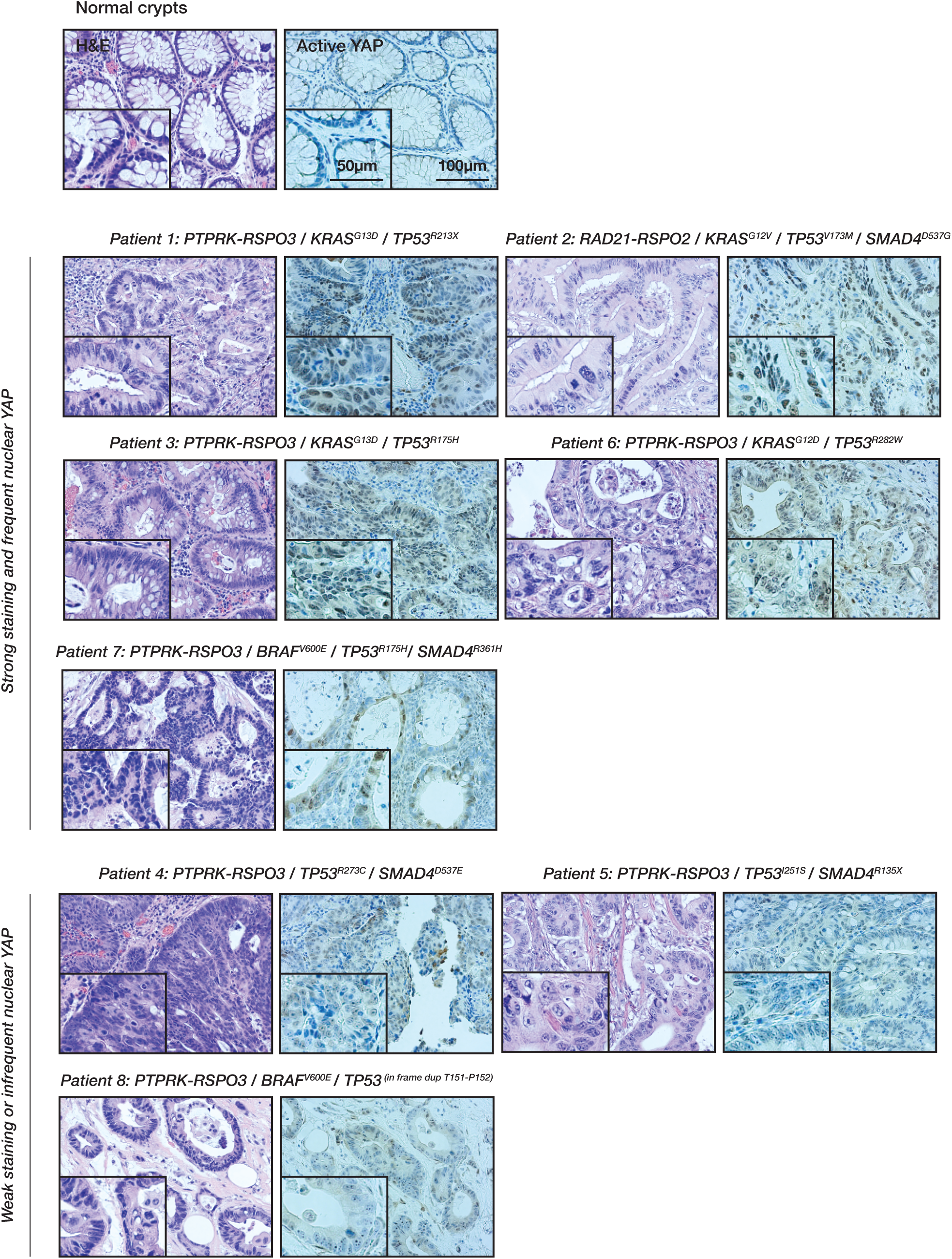
Human RSPO2/3 fusion CRCs show elevated nuclear YAP. H&E and immunohistochemical stains for active (non-phosphorylated YAP1) in 8 RSPO2 or RSPO3 fusion CRCs. Tumors were classified as those having strong and/or frequent nuclear YAP signal, or those with weak and/or infrequent YAP staining.

**Supplementary Figure 11.**
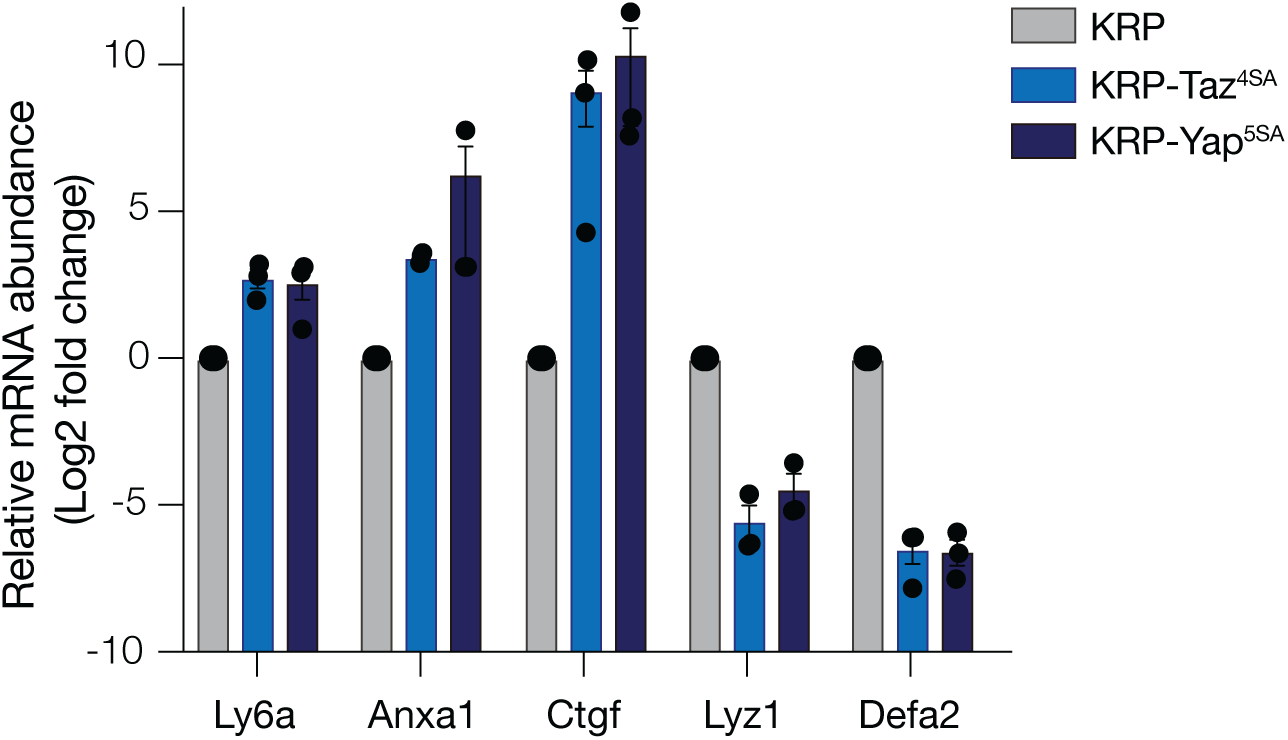
Yap/Taz promotes lineage reversion and WNT independence. qRT-PCR analysis of gene expression in KRP, KRP-TAZ^S4A^ and KRP-YAP^S5A^ organoids showing induction of YAP/TAZ targets (Ctgf) and fetal markers (Anxa1, Ly6a), and suppression of Paneth cell markers (Lyz1, Defa2). Error bars are +/- SEM.

**Supplementary Table 1:**
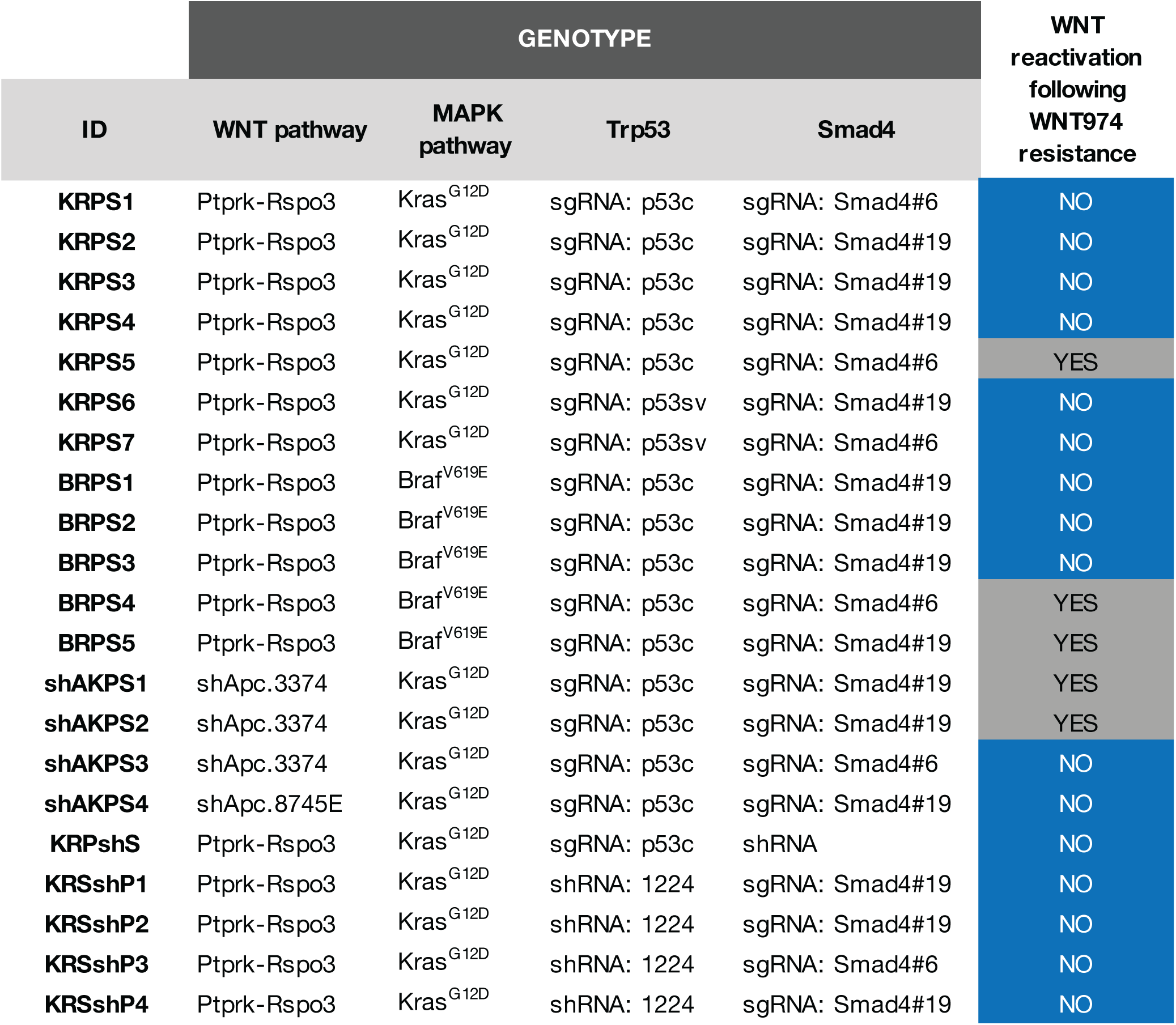
Organoid genotypes and summary of WNT974 resistance mechanisms.

